# Universal nucleic acid preservation in biological fluids with boron clusters

**DOI:** 10.64898/2026.01.13.697153

**Authors:** Brian Pak Shing Pang, Lichun Zhang, Ben Tin Yan Wong, Mike P. Williamson, Hei Ming Lai

## Abstract

Nucleic acid preservation remains a critical bottleneck for diagnostics and therapeutics, with small molecule inhibitors such as EDTA showing limited spectrum against diverse nucleases, while protein-based alternatives requiring costly cold-chain storage. Here, we report dodecaborate cluster [B_12_H_12_]^2-^ as a novel class of pan-nuclease inhibitor with a fundamentally different mechanism—preventing protein-nucleic acid interactions rather than targeting cofactor dependencies. [B_12_H_12_]^2-^ inhibited all six DNases and ten RNases tested, making it the broadest spectrum nuclease inhibitor known. Remarkably, its inhibition is reversible via γ-cyclodextrin complexation. We demonstrate that [B_12_H_12_]^2-^ preserved a physiological range of DNA for 14 days and that of RNA for 3-7 days at room temperature in human plasma and urine, achieving up to 323,972-fold better RNA retention than controls. A newly developed blood collection tube using [B_12_H_12_]^2-^ enables whole-blood circulating cell-free RNA sequencing after 7 days of room-temperature storage, with preserved transcript integrity. Mechanistic studies suggest [B_12_H_12_]^2-^ binds to nuclease active sites and creates electrostatic barriers that prevent substrate binding. This chemically stable, indefinitely shelf-stable inhibitor enables cold-chain-free biological sample transport, potentially transforming accessibility of nucleic acid-based diagnostics worldwide.

**Graphical abstract:** 

## Introduction

Nucleic acid-based technologies are foundational for modern science^[1–5]^, serving as versatile biomolecular analytes and biotechnological tools. Remarkably, nucleic acid-based technologies continue to evolve, with applications such as gene editing^[6–8]^, mRNA vaccines^[9–13]^, liquid biopsies^[14–18]^, metagenomics^[19–21]^, aptamers^[22–25]^, and spatial transcriptomics^[26–29]^ promising transformative benefits to humanity. This is a celebrated result of enabling scientific advances made in the past decades, including genetics^[30,31]^, phosphoramidite chemistry^[32,33]^, polymerase chain reaction ^[34]^, sequencing^[35–37]^, bioinformatics^[38]^, and non-inflammatory oligonucleotide modifications^[39]^.

Despite numerous scientific innovations in nucleic acid applications and methods throughout their life cycles, methods for preserving nucleic acids have remained relatively stagnant. Preserving nucleic acid integrity is crucial for the reliability of analytics, diagnostics, and therapeutics. Among the multiple nucleic acid stabilisers developed, the biochemically inert chelating agent ethylenediaminetetraacetic acid (EDTA)^[40]^ remains the preservative of choice, on the assumption that nucleic acids are only degradable in the presence of multivalent cations. However, an examination of all known nucleases reveals that this is not the case (**Figure 1a, Supp Fig 1**). Developing a nuclease inhibitor with an even broader inhibitory scope - even for nucleases yet to be discovered - must therefore act with a more general mechanism than targeting cofactor dependence. Preserving RNAs is especially problematic^[41,42]^. Many endogenous and environmental RNases, such as RNase A, are not dependent on divalent cations for their activity^[43]^. The development of RNA-based technologies, therefore, faces significantly more challenges than their DNA counterparts, which is unfortunate as the two complement each other in their utility. For example, RNA-based monitoring in biological fluids provides a dynamic view of body states^[44,45]^, compared to the static, mutational-signature-based DNA-based liquid biopsies. Meanwhile, mRNA vaccines and RNA editing do not pose a risk of genomic integration or alterations^[46,47]^.

**Figure 1.**
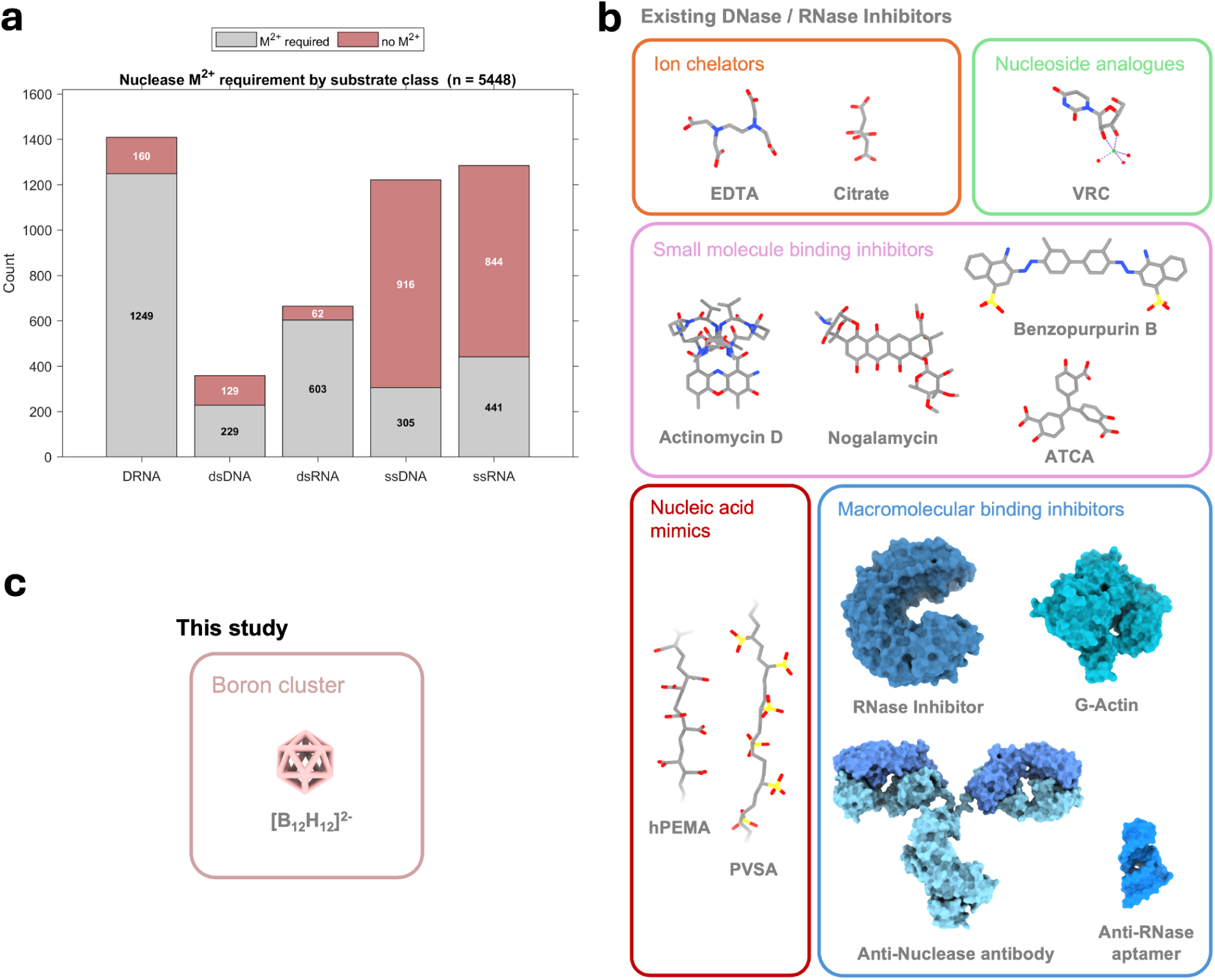
[B_12_H_12_]^2-^ as a member of a new class of nuclease inhibitor. **a.** Analysis of all known nucleases and their divalent metal ion dependence, and hence susceptibility to inhibition by ion chelators - the most widely used nuclease inhibitor. DRNA: DNA-RNA hybrid. **b.** A collection of known nuclease inhibitors and their molecular structures. ATCA: aurin tricarboxylic acid, EDTA: ethylenediaminetetraacetic acid, hPEMA: hydrolyzed poly(ethene-alt-maleic acid), PVSA: polyvinylsulfonic acid, VRC: vanadyl ribonucleoside complex, shown is an example where the base is a uridine. **c**. Structure of boron cluster as a pan-nuclease inhibitor.

Hence, the development of broad-spectrum nuclease inhibitors that minimise perturbation of bystanding biochemical processes is essential to enable continued advancements in nucleic acid-based technologies. Existing DNase and RNase inhibitors suffer from several limitations and belong to a restricted class of compounds, namely ion chelators, analogue inhibitors, and small-molecule and macromolecule antagonists (**Figure 1**)^[48]^. EDTA and citrate are the most commonly used nuclease inhibitors in blood collection tubes^[49]^. Still, they inhibit only divalent-ion-dependent nucleases due to their reliance on chelation^[50]^. Notably, EDTA is also ineffective or even stimulates certain RNase activities^[51]^. Small-molecule inhibitors are enzyme antagonists that bind to nucleases or the nucleic acid substrates. These include RNase inhibitors, such as vanadyl ribonucleoside complex (VRC)^[52]^, aurintricarboxylic acid (ATCA)^[53]^, benzopurpurin B and its derivatives^[54]^, and DNase inhibitors, including 2-nitro-5-thiocyanatobenzoic acid (NTCB)^[55]^, nogalamycin^[56]^, actinomycin D, and crystal violet^[57]^. Their scope of nuclease inhibition is relatively limited and can impact various downstream DNA and RNA analyses, including reverse transcription and sequencing. Removing any residual nuclease inhibitors cleanly proved difficult. Anionic polyelectrolytes, such as hydrolysed polyethylene maleic anhydride (hPEMA) for DNases^[55]^ and polyvinyl sulfonic acid (PVSA) for RNases^[58]^, broadly inhibit nucleases as mimics of their substrates. Still, their chemical activities are unpredictable regarding inhibitory action, selective removal, and interference with downstream assays.

Protein-based RNase and DNase inhibitors are affinity-based reagents that bind to nucleases in a 1:1 noncovalent complex, which can be naturally occurring, as with the case for actin for DNase^[59]^ and Placental RNase inhibitor for RNases ^[60]^. They cause minimal interference in subsequent assay performance and are selectively removable. However, protein-based nuclease inhibitors are costly due to their monovalency, have a short shelf life at room temperature, and are sensitive to denaturation. Even more expensive formulations consist of a mixture of anti-RNase antibodies^[61]^ and evolved mutants of thermostable RNase inhibitors, which are available commercially (such as the Superase•In RNase inhibitor from Invitrogen); nonetheless, they are still sensitive to environmental degradation in complex biological fluids due to their proteinaceous nature, and can only selectively inhibit specific nucleases by design (**Table 1**). Moreover, such inhibition is irreversible. Hence, these reagents are only laboratory-friendly but not suitable for pre-analytical nucleic acid stabilisation during transport and storage. A recent patent suggests aptamers as a replacement for antibodies, but they are not yet widely available^[62]^. To date, no studies have demonstrated selective reversal or removal of nuclease-inhibitory compounds.

**Table 1.** RNases inhibited by various RNase inhibitors. - indicates no data available.

|  | VRC | ATCA | Murine RNase Inhibitor | SUPERase-In™ | [B<sub>12</sub>H<sub>12</sub>]<sup>2-</sup> |
| --- | --- | --- | --- | --- | --- |
| RNase A | Yes | Yes | Yes | Yes | Yes |
| RNase B | - | - | Yes | Yes | Yes |
| RNases C | - | - | Yes | Yes | Yes |
| RNase I | - | - | No | Yes | Yes |
| RNase 4 | - | - | - | - | Yes |
| RNase T1 | Yes | - | No | Yes | Yes |
| RNase T2 | - |  | No | - | Yes |
| RNase U1 | - | - | No | No | - |
| RNase U2 | Yes | - | No | No | Yes |
| RNase CL1 | - | - | No | No | - |
| Nuclease P1 | No | Yes | No | - | Yes |
| RNase H | No | - | No | - | Yes |

Here, we introduce the dodecaborate cluster ion [B_12_H_12_]^2-^, as a novel broad-spectrum nuclease inhibitor (**Figure 1**). Previous studies have demonstrated that [B_12_H_12_]^2-^ allow universal inhibition of arbitrary pairs of antibody-antigen interactions^[63]^, or more generally, any protein-protein interactions without protein denaturation^[64]^ Such finding has since been applied to engineer macromolecular transport phenomena for deep and uniform three-dimensional immunohistochemistry^[63]^ and to preserve colloidal stability^[64]^. Extending the recently proposed logic of preventing biomacromolecular interactions that precede degradative reactions^[65]^, we wondered whether the same general inhibition of intermacromolecular interactions applies to interactions between proteins and nucleic acids and to the consequential nucleic acid hydrolysis catalysed by nucleases. With such a unique mode of action, [B_12_H_12_]^2-^ may broadly inhibit nucleases without requiring any prior knowledge, such as their divalent ion cofactor dependencies. In addition, [B_12_H_12_]^2-^ is exceptionally inert and can be bioorthogonally removed with size-matched supramolecular hosts, such as γ-cyclodextrin (γCD) derivatives^[66–68]^, thereby minimising interference with downstream assays.

## Results

### [B_12_H_12_]^2-^ inhibit diverse classes of deoxyribonucleases (DNases)

We began by exploring the prospects of inhibiting DNase-DNA interactions and their consequential degradation. For each DNA exconuclease and endonuclease, the nucleases and their suitable substrates were incubated with different concentrations of Na_2_[B_12_H_12_], compared with NaCl as a control. For DNase II, which is sensitive to the concentrations of certain cations, we tested and selected cations that had minimal effect on nuclease activity during the comparisons. Compared with a two-times NaCl concentration, as shown in Figure 2 and **Supp** Fig 2 **-7**, the corresponding Na_2_[B_12_H_12_] groups showed much less DNA digestion by their corresponding nuclease, implying that [B_12_H_12_]^2-^ is the key factor to inhibit DNase-DNA digestion.

**Figure 2.**
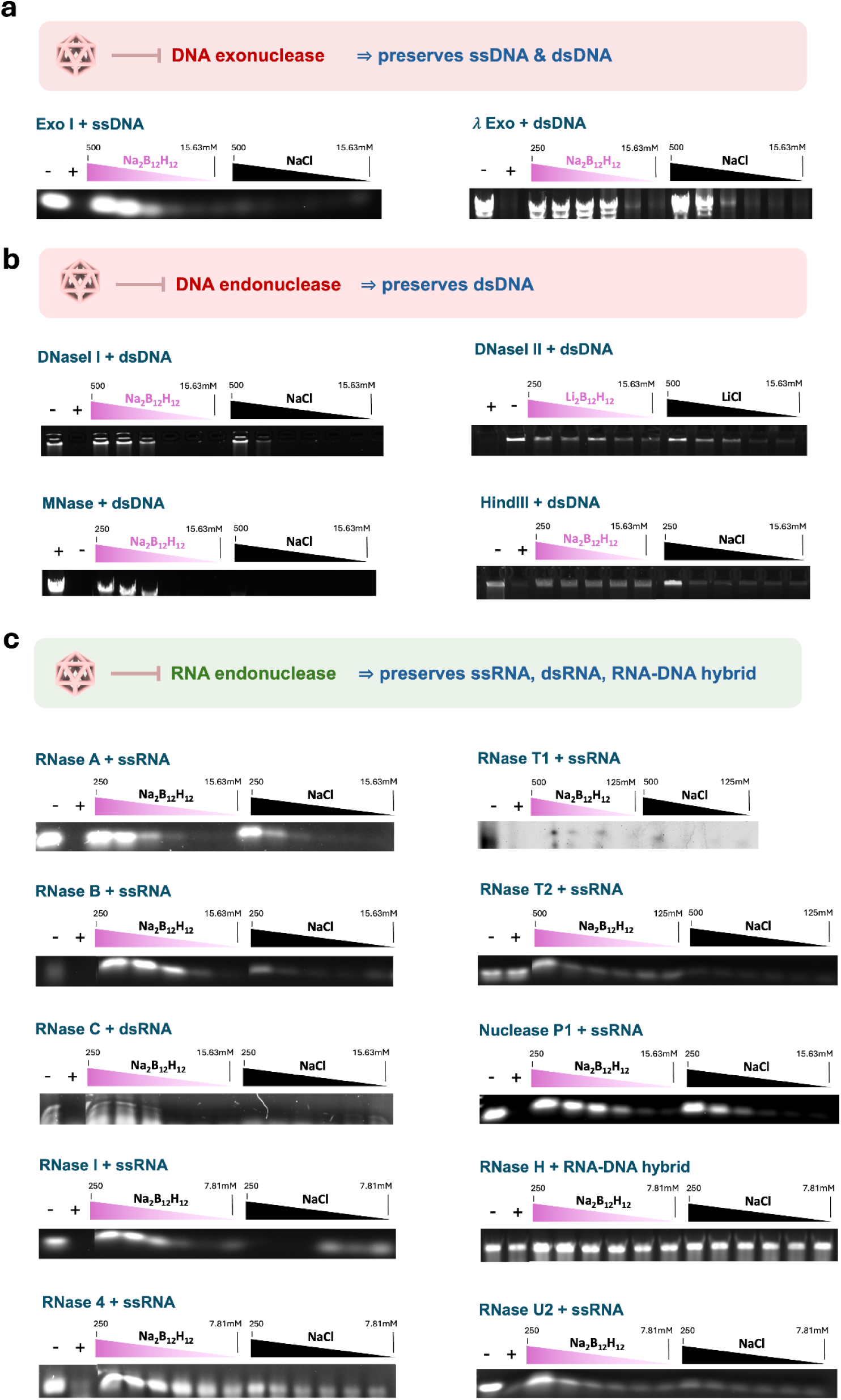
Pan-inhibition of DNases and RNases by [B_12_H_12_]^2-^. All dilution steps were two-fold; the dilution starting and ending concentrations were as labelled. + and - indicates controls with added and omitted nuclease, respectively. **a** DNA exonucleases tested. **b** DNA endonucleases tested. **c** RNA endonucleases tested.

These results suggest that [B_12_H_12_]^2-^ inhibited all DNases tested. Notably, DNase II is a lysosomal DNase that does not require a divalent cation (e.g., Ca^2+^) for function^[69]^, suggesting that [B_12_H_12_]^2-^ can be a more universal inhibitor of DNases, thereby preserving DNA in biological fluids and tissue samples. Indeed, it has been found that EDTA or EGTA can enhance nuclease activity independent of their cation chelation properties^[51]^.

### [B_12_H_12_]^2-^ inhibit diverse classes of ribonucleases (RNases)

Next, we investigated if preserving RNA is practical with such a fundamentally different inhibitory mechanism of protein-nucleic acid interaction by [B_12_H_12_]^2-^, since RNAs are sensitive to the trace amount of environmental contaminants and molecularly robust RNases.

Analogous to our DNase digestion experiments, we performed digestion experiments with various RNases on suitable RNA substrates, comparing the results between [B_12_H_12_]^2-^ salts and the corresponding Cl^-^ salts. The [B_12_H_12_]^2-^ groups also showed greater RNase inhibitory activity than the corresponding 2x Cl^-^ group controls, as shown in Figure 2c. These effects were demonstrated for all common RNases tested, all are endoribonucleases, including RNase A, RNase B, RNase I, RNase 4, RNase T2, RNase U2, Nuclease P1, digesting single-stranded yeast total RNAs (ssRNAs); RNase C (also known as RNase III), digesting double-stranded RNAs (dsRNAs); and RNase H, digesting RNA-DNA hybrids. All uncropped gel images were provided in Supp Fig 8 - 17. In our hands, RNase T1 exhibited exceptionally high activity and required 0.5 M of [B_12_H_12_]^2-^ for significant inhibition. The high concentration of [B_12_H_12_]^2-^ distorted gel assay result. With the addition of 1 U/μL SUPERase·In™, which is marketed to inhibit 0.0075 U/μL of RNase T1 for 4 hours at 37°C, most of the RNAs are still digested unless high concentrations of [B_12_H_12_]^2-^ are present. Interestingly, the inhibition of RNase seems to synergise with SUPERase·In™ in the inhibition of RNases, as shown in Supp Fig 18.

Compared with EDTA, VRC, ATCA, murine RNase inhibitor, and SUPERase·In™ - an augmented RNase inhibitor featuring the broadest RNase inhibition known, [B_12_H_12_]^2-^ exhibits the widest spectrum of inhibition, and is the only agent capable of inhibiting all the tested RNases (**Table 1**).

### Nuclease activity inhibition by [B_12_H_12_]^2-^ can be selectively removed or reinstated

Since [B_12_H_12_]^2-^ inhibit the interactions between nucleic acid substrates and the nuclease protein molecules, it is possible to reverse the inhibitory effects of [B_12_H_12_]^2-^by bio-orthogonal host-guest complexation using γ-cyclodextrin (γCD)^[66–68]^. In this study, we especially used the more water-soluble 2-hydroxypropylated (2HPγCD) derivative^[63,65]^ as illustrated in Figure 3a. The reverse of [B_12_H_12_]^2-^ inhibition by adding 2HPγCD is shown for DNase II and RNase I in Figure 3b**, c**. We subsequently confirmed the reinstatement of DNA/RNA digestion in other nucleases tested as aforementioned, by adding equimolar 2HPγCD to a given concentration of [B_12_H_12_]^2-^, as shown in Supp Fig 2 **- 7**, right-sided panels for DNases, and in Supp Fig 8 **- 17,** right-sided panels for RNases.

**Figure 3.**
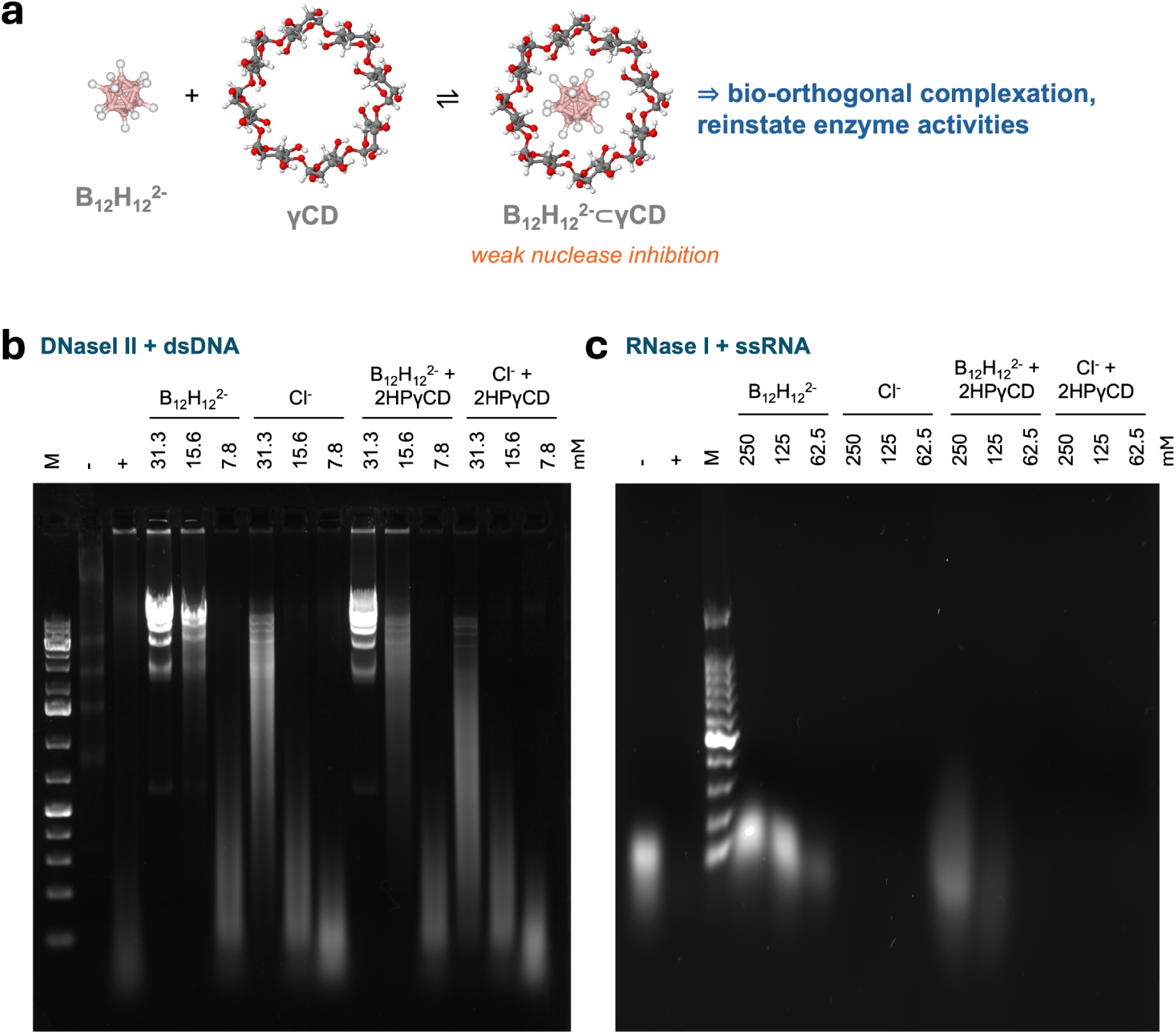
Supramolecular reversal of [B_12_H_12_]^2-^-mediated nuclease inhibition. **a** Reaction schematic of the host-guest complexation reaction involved. **b**. Reversing [B_12_H_12_]^2-^-mediated DNase II inhibition by adding 2-hydroxypropyl-γ-cyclodextrin (2HPγCD) in a 1:1 molar ratio to [B_12_H_12_]^2-^. **c**. Reversing [B_12_H_12_]^2-^-mediated RNase I inhibition by adding 2HPγCD in a 1:1 molar ratio to [B_12_H_12_]^2-^. This figure is a replica of Supp **Fig 11c**.

Hence, the [B_12_H_12_]^2-^/2HPγCD system provides an orthogonal approach to the bidirectional control of nucleic acid digestion by nucleases. The existence of an affordable, specific reversal agent makes [B_12_H_12_]^2-^ unique among the nuclease inhibitors (Figure 1).

### Preservation of nucleic acid integrity in biological fluids at room temperature

To reinforce [B_12_H_12_]^2-^ as a broad-spectrum nuclease inhibitor, we next test whether [B_12_H_12_]^2-^can preserve nucleic acids in various biological fluids. We spiked λDNA into freshly drawn human plasma at multiple dilutions with standard EDTA anticoagulation, where 0.25 M [B_12_H_12_]^2-^ was added to the test group for augmented preservation. We then incubated the plasma at room temperature for up to 14 days to simulate sample transport logistics, as illustrated in Figure 4a. Using quantitative PCR (qPCR), the EDTA-only control group showed an upward trend of cycle threshold (Ct) values from day 0 to day 14 of incubation, suggesting a continuous DNA degradation. Intriguingly, at lower concentrations of spiked-in λDNAs, both groups exhibited downward trends in Ct values over time, which was more pronounced when [B_12_H_12_]^2-^was added. This suggests that more nucleic acids are being detected over time, surpassing the value in both day 0 groups, which should be minimal in DNA degradation (Figure 4b**, Supp Fig 19a**). We hypothesised that plasma protein-binding DNA might hinder efficient denaturation and amplification of nucleic acids, and that [B_12_H_12_]^2-^ can dissociate protein-DNA complexes. Indeed, a proteinase K digestion of plasma yielded the lowest Ct values, consistent with our postulation that [B_12_H_12_]^2-^ protects DNA from degradative processes and promotes its release from plasma proteins (Supp Fig 20).

**Figure 4.**
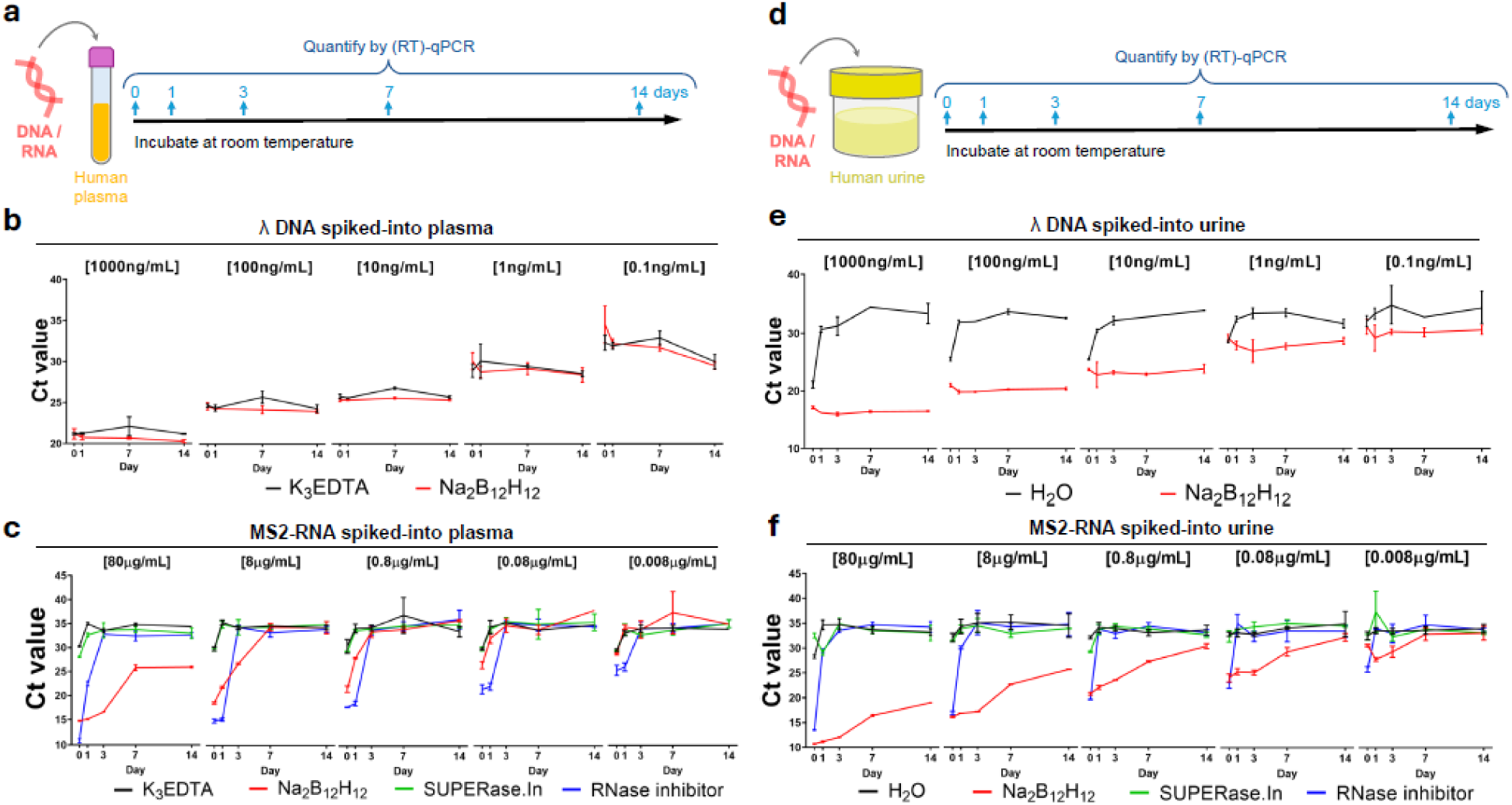
[B_12_H_12_]^2-^ preserves λ DNA and MS2 RNA in human biological fluids at room temperature. **a** Human plasma experiment schematic. **b** Plasma λ DNA levels were measured at various time-points (0, 1, 7 and 14 days) by real-time PCR (n=3). Final plasma concentration of [B_12_H_12_]^2-^ is 0.25 M, and the initial λ DNA spike-in concentration in plasma is as indicated. Bars represent group mean ± SD. **c** Plasma MS2 ssRNA levels were measured at various time-points (0, 1, 3, 7 and 14 days) by real-time PCR (n=3). Final plasma concentrations of [B_12_H_12_]^2-^, SUPERase.In and RNase inhibitor are 0.25 M, 2U/μL and 4U/μL, respectively. The initial MS2 ssRNA spike-in concentration in plasma is as indicated. Bars represent group mean ± SD. **d** Human urine experiment schematic. **e** Urine λ DNA levels were measured at various time-points (0, 1, 3, 7 and 14 days) by real-time PCR (n=3). Final urine concentration of [B_12_H_12_]^2-^ is 0.25 M, and the initial λ DNA spike-in concentration in urine is as indicated. Bars represent group mean ± SD. **f** Urine MS2 ssRNA levels were measured at various time-points (0, 1, 3, 7 and 14 days) by real-time PCR (n=3). Final urine concentrations of [B_12_H_12_]^2-^, SUPERase.In and RNase inhibitor are 0.25 M, 2U/μL and 4U/μL, respectively. The initial MS2 ssRNA spike-in concentration in urine is as indicated. Bars represent group mean ± SD.

The ability of [B_12_H_12_]^2-^ in preserving spiked-in RNA in human fluids is also evaluated. We spiked MS2 phage ssRNA into human plasma at physiologically relevant concentrations (ranging from 80 ng/mL to 80 μg/mL), incubated at room temperature for up to 7 days. Using reverse transcription-quantitative PCR (RT-qPCR), the control group showed rapid degradation of MS2 ssRNA within 30 minutes, resulting in Ct values >30. However, the Ct value of the [B_12_H_12_]^2-^ group was 15.54, suggesting 2^30-15.54^ ≈ 22,537× better retention of RNA in plasma. We also benchmarked the performance of [B_12_H_12_]^2-^ against two commercially available, protein-based RNase inhibitors. Although the recombinant murine RNase inhibitor performed the best on day 0, persistent RNase inhibition was only observed in the [B_12_H_12_]^2-^ group. [B_12_H_12_]^2-^ started out-performing the murine RNase inhibitor from day 3 (Figure 4c**, Supp Fig 19b**). This is likely due to denaturation and proteolytic degradation of the protein-based RNase inhibitor, leading to a loss of activity over time. This highlights the main disadvantages of murine RNase inhibitors and their derivatives: the protein-based inhibitors require cold-chain transport to maintain their activity, making their routine use in blood collection tubes costly and impractical. In contrast, [B_12_H_12_]^2-^ is stable indefinitely at room temperature.

Urine is a more easily accessible biological fluid for biomolecule testing than plasma, yet comes along with a challenge of higher nuclease activity. Hence, we extended our investigation into the stability of spiked-in DNA and RNA in human urine, as illustrated in Figure 4d. Repeating a similar experiment as described for human plasma, we found that [B_12_H_12_]^2-^ also led to better preservation of both DNA and RNA at all time points investigated. DNA was preserved up to 14 days at room temperature as shown in Figure 4e and Supp **Fig 21a**. Meanwhile, RNA was preserved up to 3 days at room temperature and with a slowed degradation up to 14 days of storage, as shown in Figure 4f and Supp **Fig 21b**. Compared with plasma results, the stability of RNA is noninferior to that of the protein-based murine RNase inhibitor starting from day 0.

Quantitatively, after 1 day of incubation at room temperature, [B_12_H_12_]^2-^ preserved 2^4.2^ = 18.25× more DNA for an initial spiked-in amount of 0.1 ng/mL DNA, and up to 2^14.49^ = 23170× more DNA for an initial concentration of 1000 ng/mL. The [B_12_H_12_]^2-^-mediated DNA preservation effect lasted up to 14 days at room temperature, providing excellent long-term stability. Whereas for RNA, compared to the best performing RNase inhibitor, [B_12_H_12_]^2-^ preserved 2^6.19^ = 73.25× more RNA for an initial spiked-in RNA concentration of 8 ng/mL, and up to 2^18.31^ = 323972× more RNA for an initial concentration of 80 μg/mL after 1 day of incubation at room temperature. Finally, [B_12_H_12_]^2-^ further maintained the stability of RNAs for 3 days at room temperature across all spiked-in concentrations tested.

### Adding [B_12_H_12_]^2-^ augments existing nucleic acid preservatives

We next combined 0.25M [B_12_H_12_]^2-^ with EDTA and biological fluid collection devices from Streck, and similarly quantified spiked-in experiments using qPCR/RT-qPCR. For DNA, all groups with additional [B_12_H_12_]^2-^ showed significantly lower Ct values (Supp Fig 22**, 23**), and only in plasma, a universal downward trend in Ct after 7 days of incubation at room temperature was observed (Supp Fig 22). The results for RNA are even more dramatic with spike-in and immediate quantification at day 0, where spiked-in RNAs are immediately degraded with EDTA or commercially available preservative from Streck (Supp Fig 24**, 25**). Hence, only the day 0 time point was assayed. When [B_12_H_12_]^2-^ was added, the Ct values showed an expected stepwise increase with exponentially decreasing spiked-in RNA amounts, suggesting preservation was relatively quantitative in both plasma and urine (Supp Fig 24**, 25**).

### [B_12_H_12_]^2-^ does not hinder, or even enhances, the extraction efficiencies of nucleic acids

Since nucleic acid extraction is crucial for downstream analytical processing, we tested whether it could be affected by the presence of [B_12_H_12_]^2-^ in biological fluid samples. Nucleic acids from human plasma samples were extracted immediately after spiking-in with known amounts of DNA and RNA, without the addition of γCD. Notably, the extraction of DNA from plasma was unaffected by the presence of [B_12_H_12_]^2-^ up to 0.25 M, as shown in Supp **Fig 26a**. In contrast, RNA extraction efficiency was enhanced when [B_12_H_12_]^2-^ was present as shown in Supp **Fig 26b**. The increased extraction efficiency was observed across both the phenol-chloroform method and the spin-column-based method.

### [B_12_H_12_]^2-^ preserves endogenous nucleic acid fragment sizes and RNAs for circulating cell-free RNA (ccfRNA) sequencing

As a preliminary test of the applicability of diagnostic devices, we next assessed whether [B_12_H_12_]^2-^ can help preserve endogenous nucleic acids in biological fluids. This will complement the existing approaches aimed at protecting compartment-specific nucleic acid sequence information (Figure 5a). The DNA fragment size distribution of circulating cell-free (ccf) DNA is known to correlate with various disease states, a field of study known as fragmentomics^[14,70–72]^. These DNA fragments are generated by cellular and humoral processes, which are subsequently released into the blood and are collectable for analysis. Circulating plasma nucleases, such as DNase 1 and DNase 1L3^[73]^continue to fragment nucleic acid analytes *ex vivo*, leading to pre-analytical errors. We collected fresh human plasma and compared the evolution of the nucleic acid size distribution after 7 days of room-temperature incubation in the absence or presence of [B_12_H_12_]^2-^. The capillary electrophoresis-based size distribution remained largely similar while that of the control group showed significant changes, as shown in Supp Fig 27, confirming that nucleic acid fragment sizes are better preserved when [B_12_H_12_]^2-^ is added.

**Figure 5.**
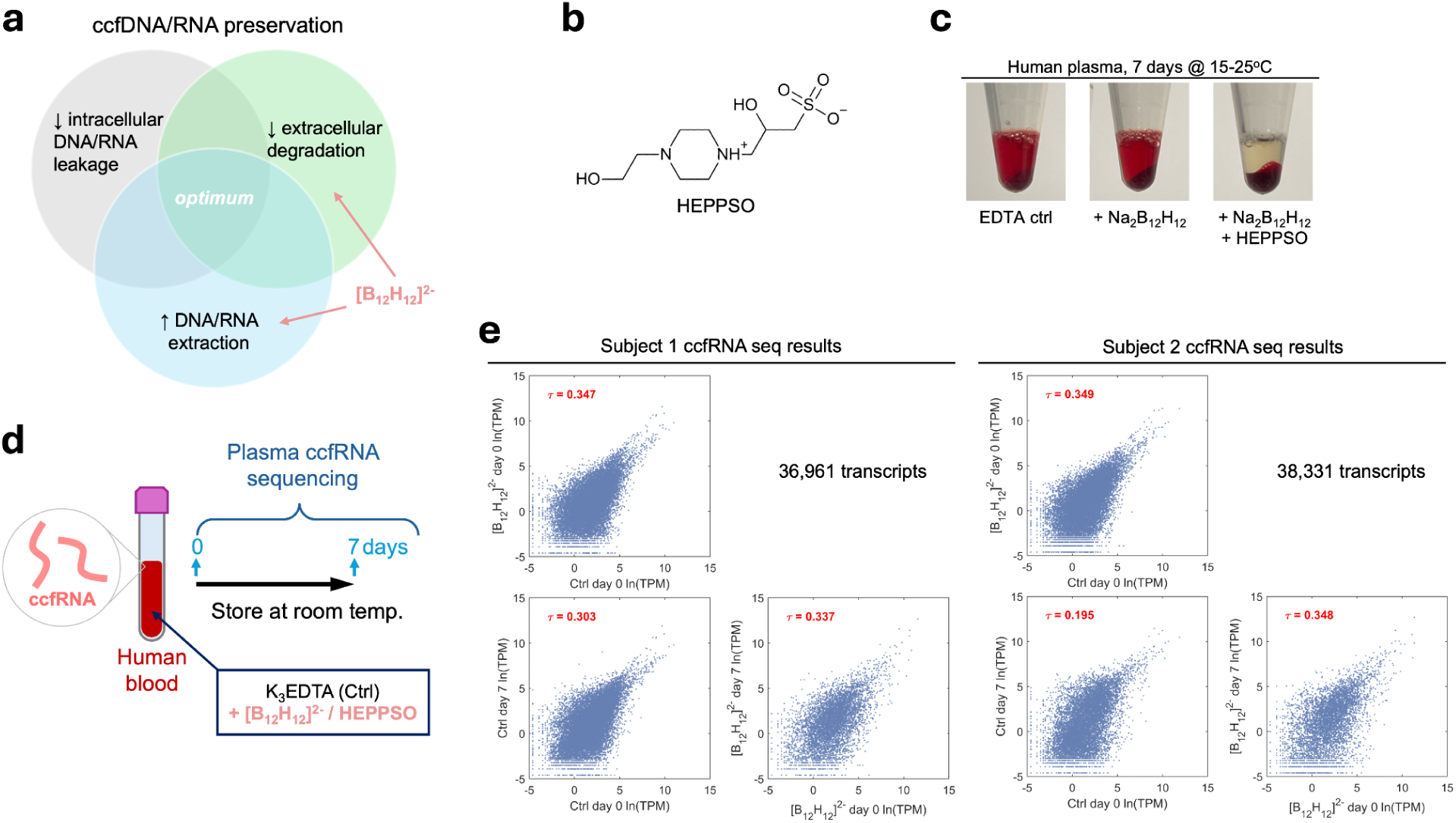
[B_12_H_12_]^2-^ improves ccfRNA sequencing reliability. **a** Summary of approaches to stabilise ccfDNA and ccfRNA analytes, and how [B_12_H_12_]^2-^ can help. **b** Chemical structure of HEPPSO. **c** Effect of HEPPSO on preventing hemolysis after prolonged room temperature storage of human whole blood. **d** Schematic of ccfRNA sequencing experiment comparing the effects of adding [B_12_H_12_]^2-^ / HEPPSO versus K_3_EDTA only control. **e** ccfRNA sequencing results for 2 subjects (a total of eight sequenced samples). The 0-day and 7-day correlation plots based on the natural log-transformed Transcripts Per Kilobase Million (ln(TPM)) were shown. τ: Kendall rank correlation coefficient.

We next tested whether [B_12_H_12_]^2-^ could preserve ccfRNAs for sequencing analysis in freshly drawn whole-blood samples. Although Na_2_B_12_H_12_ does not interact with conventional K_3_EDTA blood-collection tube components, it promotes hemolysis and thereby affects ccf-nucleic acid analytical results. To address this issue, we screened zwitterionic compounds that can be membrane-protective^[74,75]^. We found that certain Good’s buffer agents with a piperazinealkylsulfonate backbone can prevent natural blood hemolysis even after prolonged storage and in the presence of [B_12_H_12_]^2-^, which we verified via spectrophotometry of the supernatant plasma and direct observation of red blood cell (RBC) morphology (Supp Fig 28). We eventually arrived at 0.1M 4-(2-hydroxyethyl)piperazine-1-(2-hydroxypropane-3-sulfonic acid) (HEPPSO) as the optimal membrane protectant (Figure 5b**, c**). Surprisingly, HEPPSO alone prevented hemolysis better than saline control (Supp Fig 28). Using [B_12_H_12_]^2-^ combined with HEPPSO for RBC protection, we constructed a novel blood collection tube where [B_12_H_12_]^2-^and HEPPSO were pre-aliquoted into EDTA blood collection tubes, as illustrated in Figure 5d. We then performed a ccfRNA sequencing experiment on two freshly drawn blood samples using these new blood collection tubes. We extracted the nucleic acids on days 0 and 7 of room-temperature storage, as illustrated in Figure 5d. The pre-sequencing electropherograms for the control group showed greater changes than those of the Na_2_B_12_H_12_/HEPPSO group **(Supp** Fig 29). This suggests more substantial nucleic acid fragmentation and degradation in the control, even though some hemolysis still occurred in the Na_2_B_12_H_12_/HEPPSO group, resulting in a new 18S rRNA peak at ∼1800 bp. All sequencing experiments were successful and the number of clean, mapped reads is similar across all samples, although we observed slightly less reads in the Na_2_B_12_H_12_/HEPPSO group (Supp **Table 2**). Using transcript-focused analysis and taking each group’s day 0 transcriptome as the reference, the RNA reads of the [B_12_H_12_]^2-^-group showed higher correlation across the two time points, as shown in Figure 5e, where Kendall’s τ correlation coefficients for the two subjects were 0.337 and 0.348 for the [B_12_H_12_]^2-^-group after 7 days of storage, while those for the control group dropped to 0.303 and 0.195 after 7 days. A subset analysis shows that [B_12_H_12_]^2-^ protection is more pronounced for transcripts >1000 nt, as indicated by larger differences in the correlation coefficients between the [B_12_H_12_]^2-^ and control groups (Supp Fig 30). Such an observation is consistent with our findings from the MS2-RNA spike-in and recovery experiment described in Figure 4.

### [B_12_H_12_]^2-^ induces repulsive electrostatic interactions between nuclease and nucleic acids

To investigate the generalised mechanism behind [B_12_H_12_]^2-^-mediated nuclease inhibition, we titrated increasing concentrations of [B_12_H_12_]^2-^ and monitored the chemical shifts of all residues on the model protein barnase (an RNase from *Bacillus amyloliquefaciens)* with NMR^[76]^. By fitting a curve to each *k*-th residue’s amide proton ^1^H*_N_* chemical shifts (δ) with varying concentrations *c* of the co-solute, and decomposing the curve into a saturable Langmuir isotherm part and a linear gradient (*m_L_*) part, one can obtain the ensemble-averaged co-solute binding affinity (*K*_d_) and solvent environment fluctuation around the *k*-th residue amide proton, respectively (**Equation 1**).

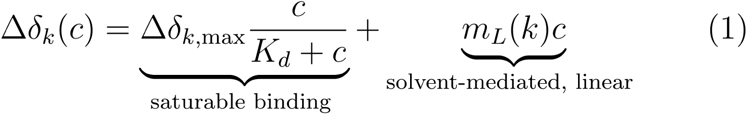

[B_12_H_12_]^2-^ showed strong interactions with many barnase residues (Fig 6a**, Supp** Fig 31), with an average *K*_d_ of 4.61 ± 2.17mM, compared to 22mM for thiocyanate and sulphate ions. This is in keeping with previous studies that boron clusters of the [B_10_X_10_]^2-^ and B_12_X_12_ ^2-^ type (X = H or halogens) can interact with albumins^[77,78]^. In addition, the per-residue *m_L_* were strongly positive, with a distribution of +263 ± 464 ppb/M, compared to –51 ± 51 ppb/M for thiocyanate and +7 ± 22 ppb/M for sulphate (Fig 6b, c)^[76]^. These results were in stark contrast to previous proposals where [B_12_H_12_]^2-^ is a weakly coordinating ion with strong chaotropic properties. Instead, the NMR titration experiment suggests that [B_12_H_12_]^2-^ interacts more strongly with proteins at specific residues compared to other common solutes, with extreme kosmotropic effects in the vicinity of macromolecules (Fig 6d). To exclude the possibility of salt contaminants, which can produce such extreme results, we performed Inductively Coupled Plasma Optical Emission Spectroscopy (ICP-OES, Supp **Table 1**) and Electrospray Ionisation-Mass Spectrometry (ESI-MS, Supp Fig 32) to confirm the absence of substantial contaminants in the commercial Na_2_B_12_H_12_ sample used that were not detectable by NMR.

**Figure 6.**
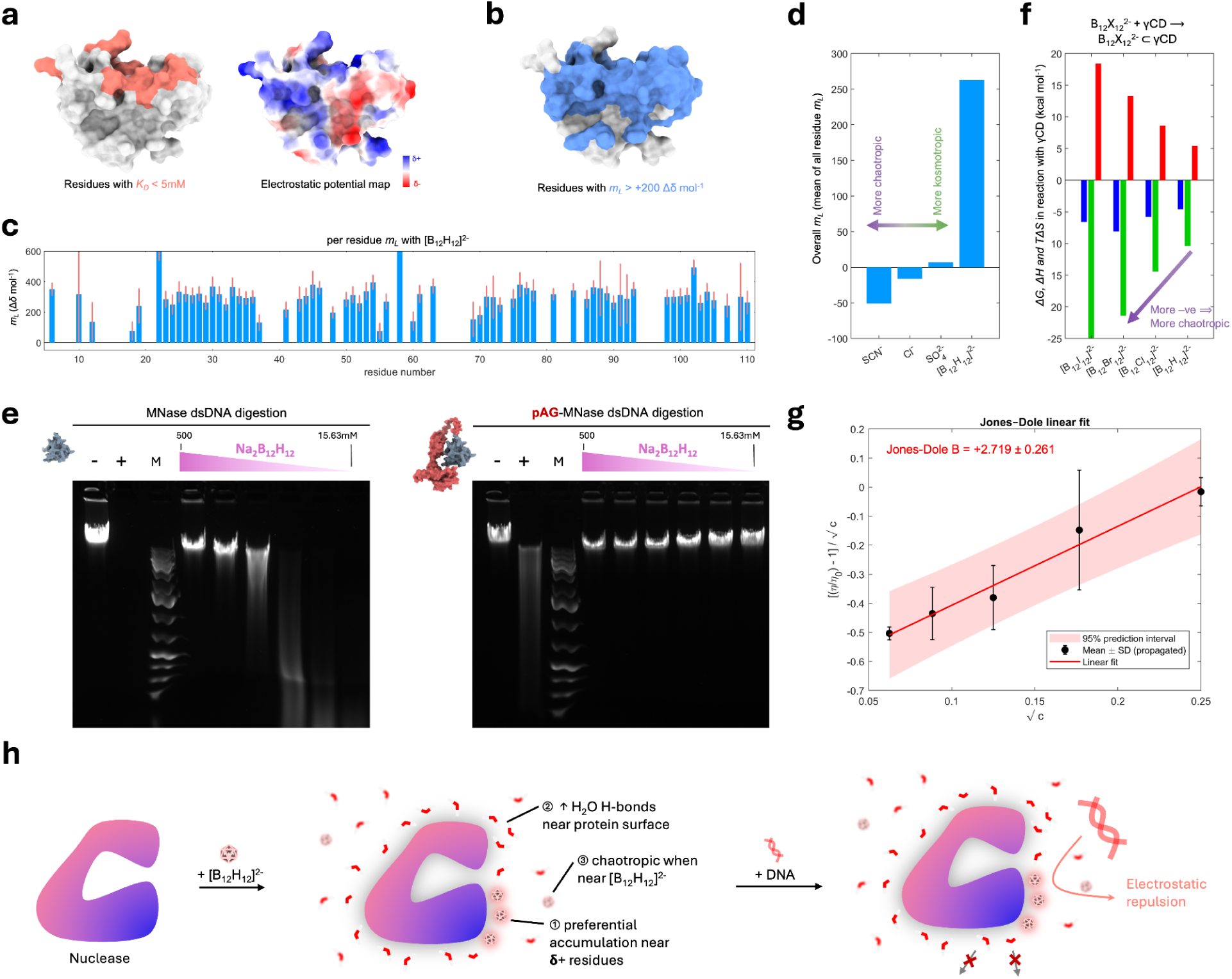
Mechanistic studies and proposed model on [B_12_H_12_]^2-^-mediated nuclease inhibition. **a** NMR titration using barnase as a model protein reveals [B_12_H_12_]^2-^ binds to residues near the active site (left), which are mostly positively charged, as seen on the electrostatic map (right). **b** NMR titration also revealed [B_12_H_12_]^2-^ drastically reduced fluctuations around the ^1^H*_N_* protons throughout the protein surface, colored in blue. **c** Plots of *m_L_* values for each residue of barnase when titrated with increasing [B_12_H_12_]^2-^ concentrations. **d** Comparison of *m_L_* values from Trevitt et al^[76]^ and those for [B_12_H_12_]^2-^. A positive (+ve) and negative (-ve) value correlates with the Hofmeister series description of kosmotropic and chaotropic ion character and is physically related to the water fluctuations near the protein residues. **e** Comparison of Δ*G*, Δ*H* and *T*Δ*S* values for the host-guest reaction between [B_12_H_12_]^2-^ and γCD in water, with other boron cluster ions’ data from Asaaf et al^[67]^. A −ve ΔH value with +ve *T*Δ*S* indicates chaotropic character of the ion being complexed. **f** Comparison of nuclease activity inhibition by [B_12_H_12_]^2-^ for MNase and pAG-MNase fusion protein, which has more surface area. Although the proteins have similar baseline activities, pAG-MNase, which has 2.8x more solvent-accessible surface area than MNase, is much more sensitive to inhibition by [B_12_H_12_]^2-^. **g** Viscosity analyses of solutions with various [B_12_H_12_]^2-^ concentrations with Jones-Dole fit. The Jones-Dole B coefficient is obtained as the slope of the fitted line. The obtained values are +2.719 ± 0.261 (standard error of fitted line slope). **h** Mechanistic model of [B_12_H_12_]^2-^-nuclease inhibition. [B_12_H_12_]^2-^ preferentially accumulates near positively charged residues (1), and reduces solvent fluctuations and hence increases H-bonds near protein surfaces (2). However, [B_12_H_12_]^2-^ reduces H-bonds in the bulk water environment far from protein surfaces (3). This leads to electrostatic repulsion of negatively charged nucleic acids from binding to nuclease active sites (1), but also makes the displacement of water molecules bound to protein surfaces unfavourable due to the excess enthalpy required to break the H-bonds between the water molecules, due to (2) and (3).

A close examination of the results suggests [B_12_H_12_]^2-^ preferentially interacts with positively charged residues (R, K), aromatic residues (Y, W, H), as well as residues capable of hydrogen bonding (Q, N) - reflective of electrostatic interactions, anion-π interactions, and hydrogen-hydride bonding (Supp Fig 31). Therefore, [B_12_H_12_]^2-^ likely binds to positively charged nuclease active sites and electrostatically repels negatively charged nucleic acid substrates, constituting its general mechanism of inhibition. Meanwhile, the water molecules around proteins are more ordered with an increased mean number of hydrogen bonds (H-bonds). Since inter-macromolecular interactions displace a large amount of water molecules near the protein surfaces, the increased number of H-bonds between the water molecules makes them thermodynamically unfavourable to be displaced - essentially forming a “solvent barrier” preventing inter-macromolecular interactions.

Unfortunately, the RNase activity of barnase was too low for reliable enzymatic studies and further biophysical studies (Supp Fig 33). To seek validation at the functional level, we conducted experiments with nucleases of varying surface areas, for which [B_12_H_12_]^2-^ can bind. Nucleases with a larger protein surface area will bind [B_12_H_12_]^2-^ readily and hence be more susceptible to its inhibition. We thus compared micrococcal nuclease (MNase) with its Protein A/G (pAG)-fused counterpart (MNase-pAG), which has 2.8x more solvent-accessible surface area, and both of which are commercially available with similar RNase activities (Fig 6e). Interestingly, RNase B - a glycosylated form of RNase A, is not more sensitive to [B_12_H_12_]^2-^inhibition (Fig 2c), suggesting that [B_12_H_12_]^2-^ has a higher affinity only to protein residues.

To understand the discrepancy with the reported chaotropic properties of [B_12_H_12_]^2-^, we further explored other biophysical properties of [B_12_H_12_]^2-^. Thermodynamic ITC analysis on the complexation of [B_12_H_12_]^2-^ by γ-cyclodextrin showed a large enthalpy contribution with entropic penalties (Fig 6f**, Supp** Fig 34), which is similar for other boron cluster ions that are also superchaotropic. We further measured changes in the relative dynamic viscosity (*η*/*η_0_*) with increasing [B_12_H_12_]^2-^ concentrations (*c*) using a Ubbelohde viscometer at 20°C. Fitting the data to the Jones-Dole equation,

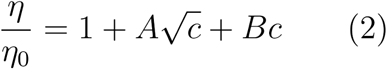

where a negative B-coefficient is suggestive, but not necessarily confirms^[79]^, the breaking of water hydrogen bonds. We found the B-coefficient for [B_12_H_12_]^2-^ to be +2.719 ± 0.261, suggestive of increased water “structuredness” (Figure 6g). However, the B-coefficient for [B_12_Cl_12_]^2-^ is +3.040 ± 0.642 further down the series, which is also positive (Supp Fig 35).

These experimental results confirm that the effects of [B_12_H_12_]^2-^ on water as a solvent are highly nuanced, defying the simplistic one-dimensional model given by the century-old Hofmeister series. While the NMR and viscosity B-coefficient provided strong evidence for [B_12_H_12_]^2-^increasing the structuredness of water in the bulk solvent, the evidence from supramolecular complexation thermodynamics is equally strong. Experimental measurements further substantiate the major role of solvent water molecules in the complexation reaction, showing that the vertical detachment energy change (ΔVDE) decreases in the order Cl > Br > I *in vacuo* ^[80]^.

Overall, our experimental evidence suggests that [B_12_H_12_]^2-^ binds tightly to nuclease active sites and reduces fluctuations in water molecules around protein surfaces and in the bulk solvent, thereby rendering enzyme-substrate binding electrostatically unfavourable (Figure 6h). We also uncovered a surprisingly complex behaviour of [B_12_H_12_]^2-^ in water, which we discuss further below.

## Discussion

We have extended the boron cluster-mediated inhibition of protein-protein interactions to protein-nucleic acid interactions. By systematically studying the effects of [B_12_H_12_]^2-^ on diverse nucleases, our study has established boron clusters as a new class of nuclease inhibitors. The unique inhibitory mechanism of shielding nucleases their nucleic acid substrates instead of targeting their cofactor dependence leads to an unmatched scope of nuclease inhibition, and hence much more robust nucleic acid protection. Owing to such a general inhibitory mechanism, chemical stability, and biological inertness, [B_12_H_12_]^2-^ is especially useful in raw biological samples where the nucleases present can remain unidentified.

Through a series of experiments, we confirmed [B_12_H_12_]^2-^ preserves nucleic acid analytes in human biological fluid samples and enhances their extraction using conventional methods. We also showed that [B_12_H_12_]^2-^ can be directly combined with existing nucleic acid preservation protocols to improve their performance, supporting its utility as a novel nucleic acid analyte preservative in real-world scenarios, especially in urine samples where nuclease activities are high^[81]^. Notably, we also demonstrated the preservation of ccfRNA in whole blood samples collected and stored in the presence of [B_12_H_12_]^2-^ for up to 7 days without refrigeration. Taken together, [B_12_H_12_]^2-^ is an accessible, immediately deployable nucleic acid preservative that augments existing approaches. The greatly stabilized DNAs and RNAs in urine samples, and likely in other biological fluids such as saliva, sputum, pleural and peritoneal fluids, cannot be understated. With the ease of urine collection, one can envision biological fluids being readily collected by patients at home and sent to specialised centres for processing without the need for cold-chain transport. Our discovery provides the foundation for universal, affordable cancer and infectious disease screening that can be cost-effectively deployed at scale. This is also likely to benefit laboratory-based workflows involving sensitive RNA analytes and to extend the shelf life of nucleic acid-based therapeutics.

Although [B_12_H_12_]^2-^ effectively inhibits many nucleases, its effects on downstream workflows, such as various sequencing technologies, are yet to be explored and will require future work. In our ccfRNA sequencing experiment, even though we observed a better correlated transcriptome after 7-day room temperature whole blood incubation, there is a slightly less overall number of reads in the [B_12_H_12_]^2-^ group, as shown in Supp **Table 2**. It is possible that some [B_12_H_12_]^2-^ may persist in the nucleic acid sample, leading to interference in downstream sequencing applications. Hence, it would be best to selectively and cleanly remove any [B_12_H_12_]^2-^ from the extracted nucleic acids. This may be achieved using γCD polymers^[82–85]^ or derivatives immobilised on solid supports^[86–88]^, such as latex or magnetic beads. In addition, even though the addition of HEPPSO prevented gross and microscopic hemolysis in the presence of [B_12_H_12_]^2-^, it is possible that analytes can traverse across the cell membrane^[89,90]^, making [B_12_H_12_]^2-^-preserved blood potentially unsuitable for other diagnostic purposes. Hence, further exploration on the compatibility of [B_12_H_12_]^2-^ with clinical assays is required.

Our mechanistic studies provided further insights into how [B_12_H_12_]^2-^ interacts with biomacromolecules^[64]^, yielding surprising findings. [B_12_H_12_]^2-^ strongly interacted with a surface patch distinct from that of previously tested ions^[76]^. This is in keeping with other reports showing that centrosymmetric boron clusters strongly adhere to proteins^[77,78]^, to the extent that controllable supramolecular assemblies can be formed^[91]^ Meanwhile, the reversibility of nuclease inhibition confirmed that the proteins are functionally intact, as previously reported ^[63,64,92]^. It is likely that the weakly coordinating character of [B_12_H_12_]^2-^ makes it thermodynamically unfavourable for it to “melt” into the backbone of proteins that would lead to their denaturation, as observed with other conventional chaotropes such as urea and guanidinium ions that directly interact with the peptide backbone. This represents a noteworthy exception in contemporary statistical mechanical theories of protein stability, where a generalised cosolute can accumulate preferentially around proteins without denaturing them^[93]^.

At the same time, we found that the effects of boron cluster ions on water fluctuations (or more traditionally, water structuredness) differ between the bulk and interfacial cases. Although NMR and B-coefficient suggest that, near proteins and in bulk water, these ions suppress water fluctuations, analyses of interfacial phenomena such as host-guest complexation and the flotation of boron clusters^[94]^ are strongly driven by their chaotropic character. As these experiments probe different aspects of the ion-solvent environment, a possible resolution is that water near boron clusters shows increased fluctuations, whereas water farther from them shows reduced fluctuations compared to pure water. In this way, [B_12_H_12_]^2-^’s behaviour defies classification by the Hofmeister series due to nuances in balancing multiple interaction energetics depending on its environment.

To conclude, we discovered boron clusters as a novel class of nuclease inhibitors with superior performance. It readily improves diagnostic performance by better preserving nucleic acid analytes, even in the presence of unknown nucleases.

## Materials and Methods

### Materials

#### Chemicals and reagents

Sodium dodecahydro-*closo*-dodecaborate ([B_12_H_12_]^2-^) was procured from Katchem (Cat. no. 257) and used without further purification. 2-hydroxypropyl-γ-cyclodextrin was obtained from Cyclolab (Cat. no. 38), Cyclodextrin Shop (Cat. no. CDexG-075), or Sigma Aldrich (Cat. no. 779229). 3-[4-(2-Hydroxyethyl)piperazin-1-yl]propane-1-sulfonic acid (EPPS) was purchased from Sigma Aldrich (cat no. E9502). HEPES was purchased from Aladdin (cat no. H109407). Trimethylamine N-oxide (TMAO) was purchased from Alfa Aesar (cat no. A14916-06). 1,4-Piperazine bis(propanesulfonic acid) (PIPES) was purchased from Alfa chemistry (cat no. ACM5625569). N,N-Bis(2-hydroxyethyl)-2-aminoethanesulfonic acid (BES, cat no. B0909), 4-(2-Hydroxyethyl)piperazine-1-(2-hydroxypropane-3-sulfonic acid) (HEPPSO, cat no. H0772) and 2-Hydroxy-4-morpholinepropanesulphonic acid (MOPSO, cat no. H0671) were purchased from TCI Chemicals.

Exonuclease I (*E. coli*) (cat no. M0293), λ exonuclease (cat no. M0262S), HindIII restriction endonuclease (cat no. R3104S), Nuclease P1 (cat no. M0660S), and Ribonuclease IV (cat no. M1284L) were purchased from New England Biolabs. Deoxyribonuclease I (cat no. EN0521), Ribonuclease A (cat no. R1253), Ribonuclease I (cat no. EN0602), Ribonuclease III (cat no. AM2290), Ribonuclease T1 (cat no. EN0541), and SUPERase·In™ RNase Inhibitor (cat no. AM2694) were purchased from Thermo Fisher Scientific. Deoxyribonuclease II was purchased from MedChemExpress (cat no. HY-P2863). Micrococcal Nuclease (cat no. LS004798), Ribonuclease T2 (cat no. LS01502), and Ribonuclease U2 (cat no. LS01520) were purchased from Worthington Biochemical. Ribonuclease B was purchased from RayBiotech (cat no. P61823). Ribonuclease H was purchased from Qiagen (cat no. Y9220L). Barnase (cat no. MBS1146186) and Barstar (cat no. MBS1053017) were purchased from MyBioSource. RNase inhibitor was purchased from Molecular Cloning Laboratories (cat no. RNIN200).

λ DNA-HindIII Digest (cat no. N3012) and dsRNA Ladder (cat no. N0363S) were purchased from New England BioLabs. λ DNA was purchased from Takara Bio (cat no. 3010). ssRNA was purchased from Millipore (cat no. 55714). tRNA (cat no. 10109495001) and MS2 single-stranded RNA genome (cat no. 10165948001) were purchased from Roche.

Proteinase K was purchased from New England Biolabs (cat no. P8107S). QIAamp ccfDNA/RNA Kit was purchased from Qiagen (cat no. 55184). Phenol/chloroform/isoamyl alcohol was purchased from Thermo Fisher Scientific (cat no. 15593031). Tripotassium ethylenediaminetetraacetate (K_3_EDTA) solution was purchased from Aladdin (cat no. E111797). Nucleic Acid BCT™ (cat no. 230644) and Streck^®^ Urine Preserve (cat no. 230599) were purchased from Streck. AGANI™ 23-G X 1.25” needle was procured from Terumo (cat no. AN*2332R1). TB Green® Premix Ex Taq™ (Tli RNaseH Plus) (cat no. RR420) and One-Step TB Green® PrimeScript™ RT-PCR Kit II (Perfect Real Time) (cat no. RR086) were purchased from Takara Bio.

Reaction buffers, including 10×NEBuffer r1.1 (cat no. B6001S), 10× NEB 2.1 buffer (cat no. B7202S), and 10× Nuclease P1 Reaction Buffer (cat no. B0660S), were purchased from New England Biolabs. 10× RNase III Reaction Buffer (cat no. AM4019G) and 20×SSC buffer (cat no. AM4019G) were purchased from Thermo Fisher Scientific. 10×RNase H buffer was purchased from Qiagen (cat no. B9220L). 50×TAE buffer was purchased from Phygene Biotechnology (cat no. LA0612). 10×TBE buffer was purchased from Bioss Antibodies (cat no. C1056). Super GelRed^TM^ was purchased from US Everbright (cat no. S2001).

## Methods

**Exo I (Exonuclease I, *E. coli*)** activity was measured using a ∼50 nt-long single-stranded DNA as the substrate. Each reaction mixture (total volume 10 μL) included 1×NEB 2.1 buffer, 1 μg of single-stranded DNA, and 10-20 U Exo I. The reactions were incubated at room temperature for 10 minutes, and then stopped by heating at 65°C for 10 minutes. The reaction products (5 μL) were analysed by electrophoresis on a 0.6% agarose gel in 1×TAE buffer, stained with Super GelRed^TM^, run at 120V for 30 minutes, and imaged using a BioRad Gel Doc EZ imager.

**λ Exonuclease** activity was measured using λ DNA-HindIII Digest as the substrate. Each reaction mixture (total volume 10 μL) included 0.5 μg of DNA substrate and 25 U λ exonuclease. The reactions were incubated at 37°C for 30 minutes, and then stopped by heating at 75°C for 10 minutes. The reaction products (5 μL) were analyzed by electrophoresis on a 0.6% agarose gel in 1×TBE buffer, stained with Super GelRed^TM^, run at 120V for 30 minutes, and imaged using a BioRad Gel Doc EZ imager.

**HindIII restriction endonuclease** activity was measured using λ DNA as the substrate. Each reaction mixture (total volume 25 μL) contained 1×NEB 2.1 buffer, 0.5 μg of the DNA substrate, and 10 U HindIII. The reactions were incubated at 37°C for 2 hours, and then stopped by heating at 65°C for 10 minutes. The reaction products (10 μL) were analyzed by electrophoresis on a 0.6% agarose gel in 1×TAE buffer, stained with Super GelRed^TM^, run at 120V for 30 minutes, and imaged using a BioRad Gel Doc EZ imager.

**DNase I (Deoxyribonuclease I)** activity was measured using λ DNA-HindIII Digest as the substrate. Each reaction mixture (total volume 10 μL) included 0.5 μg of DNA substrate and 0.5 U DNase I. The reactions were incubated at room temperature for 10 minutes, then stopped by adding 1 μL of 0.5M EDTA buffer (pH 8.0). The reaction products (5 μL) were analyzed by electrophoresis on a 0.6% agarose gel in 1×TBE buffer, stained with Super GelRed^TM^, run at 120V for 40 minutes, and imaged using a BioRad Gel Doc EZ imager.

**DNase II (Deoxyribonuclease II)** activity was measured using λ DNA-HindIII Digest as the substrate. Each reaction mixture (total volume 10 μL) included 0.5 μg of DNA substrate and 0.5 U DNase II. The reactions were incubated at room temperature for 5-15 minutes, and then stopped by heating at 65 °C for 10 minutes. The reaction products (5 μL) were analyzed by electrophoresis on a 0.6% agarose gel in 1×TAE buffer, stained with Super GelRed^TM^, run at 120V for 40 minutes, and imaged using a BioRad Gel Doc EZ imager.

**MNase (Micrococcal Nuclease)** activity was measured using λ DNA as the substrate. Each reaction mixture (total volume 10 μL) included 1×reaction buffer (5mM CaCl2, 50mM Tris-HCl, pH 8.0), 0.5 μg of DNA substrate, and 1 U MNase. The reactions were incubated at 37°C for 15 minutes, then stopped by adding 1 μL of 0.5 M EGTA buffer (pH 8.25). The reaction products (5 μL) were analyzed by electrophoresis on a 1% agarose gel in 1×TBE buffer, stained with Super GelRed^TM^, run at 120V for 30 minutes, and imaged using a BioRad Gel Doc EZ imager.

**Nuclease P1** activity was measured using ssRNA as the substrate. Each reaction mixture (total volume 10 μL) included 1×Nuclease P1 Reaction Buffer, 1 μg of RNA substrate, and 0.5 U Nuclease P1. The reactions were incubated at 37°C for 30 minutes, and then stopped by heating at 75 °C for 10 minutes. The reaction products (5 μL) were analyzed by electrophoresis on a 1% agarose gel in 1×TBE buffer, stained with Super GelRed^TM^, run at 120V for 30 minutes, and imaged using a BioRad Gel Doc EZ imager.

**RNase A (Ribonuclease A)** activity was measured using ssRNA as the substrate. Each reaction mixture (total volume 10 μL) included 0.5 μg of RNA substrate and 50 mU RNase A. The reactions were incubated at room temperature for 10 minutes, then stopped by adding excess (5 μL) RNase inhibitor. The reaction products (5 μL) were analyzed by electrophoresis on a 0.6% agarose gel in 1×TBE buffer, stained with Super GelRed^TM^, run at 120V for 30 minutes, and imaged using a BioRad Gel Doc EZ imager.

**RNase B (Ribonuclease B)** activity was measured using ssRNA as the substrate. Each reaction mixture (total volume 10 μL) included 1 μg of RNA substrate, and 0.1 U RNase B. The reactions were incubated at 37°C for 15 minutes, then stopped by adding excess RNase inhibitor (5 μL). The reaction products (5 μL) were analyzed by electrophoresis on a 1% agarose gel in 1×TBE buffer, stained with Super GelRed^TM^, run at 120V for 30 minutes, and imaged using a BioRad Gel Doc EZ imager.

**RNase I (Ribonuclease I)** activity was measured using ssRNA as the substrate. Each reaction mixture (total volume 10 μL) included 1 μg of RNA substrate, and 1 U RNase I. The reactions were incubated at room temperature for 10 minutes, and then stopped by adding 1 μL of 1% SDS. The reaction products (5 μL) were analyzed by electrophoresis on a 1% agarose gel in 1×TBE buffer, stained with Super GelRed^TM^, run at 120V for 30 minutes, and imaged using a BioRad Gel Doc EZ imager.

**RNase III (Ribonuclease III, or Ribonuclease C)** activity was measured using dsRNA Ladder as the substrate. Each reaction mixture (total volume 10 μL) included 1×RNase III Reaction Buffer, 0.1 μL of RNA substrate, and 0.1 U RNase III. The reactions were incubated at 37°C for 1 hour, then stopped by adding 1 μL of 0.5 M EDTA buffer (pH 8.0). The reaction products (5 μL) were analyzed by electrophoresis on a 1% agarose gel in 1×TBE buffer, stained with Super GelRed^TM^, run at 120V for 30 minutes, and imaged using a BioRad Gel Doc EZ imager.

**RNase IV (Ribonuclease IV)** activity was measured using denatured tRNA as the substrate. tRNA was denatured by heating in 3 M urea at 90°C for 10 minutes, cooled down at room temperature, and then diluted with nuclease-free water to a final urea concentration of 1 M. Each reaction mixture (total volume 10 μL) included 1XNEBuffer r1.1, 0.33 μg of RNA substrate, and 25 U RNase IV. The reactions were incubated at 37°C for 1 hour, then stopped by adding excess RNase inhibitor (5 μL). The reaction products (5 μL) were analyzed by electrophoresis on a 1% agarose gel in 1×TBE buffer, stained with Super GelRed^TM^, run at 120V for 30 minutes, and imaged using a BioRad Gel Doc EZ imager.

**RNase H (Ribonuclease H)** activity was measured using RNA-DNA hybrid as the substrate. RNA-DNA hybrid was generated by incubating MS2 single-stranded RNA genome with a complementary DNA oligo (5’-TGATGGACTCACCC-3’) in 1 to 5 molarity ratio in 1×SSC buffer. The nucleic acid mixture was denatured at 95°C for 5 minutes, then cooled to room temperature at a rate of −0.1°C per second. Each RNase H reaction mixture (total volume 10 μL) included 1×RNase H buffer, ∼0.3 μg of freshly prepared RNA-DNA hybrid substrate, and 5 U RNase H. The reactions were incubated at 37°C for 1 hour, then stopped by adding 1 μL of 0.5 M EDTA buffer (pH 8.0). The reaction products (5 μL) were analyzed by electrophoresis on a 1% agarose gel in 1×TBE buffer, stained with Super GelRed^TM^, run at 120V for 30 minutes, and imaged using a BioRad Gel Doc EZ imager.

**RNase T1 (Ribonuclease T1)** activity was measured using ssRNA as the substrate. Each reaction mixture (total volume 10 μL) included 1 μg of RNA substrate and 1 U RNase T1. SUPERase•In^TM^ RNase Inhibitor at 2 U/uL was used in the control for benchmarking. The reactions were incubated at 37°C for 4 hours, then quenched by adding 1 μL of 1% SDS. The reaction products (10 μL) were analyzed by electrophoresis on a 10% Urea-PAGE gel in 1×TBE buffer, stained with Super GelRed^TM^, run at 120V for 45 minutes, and imaged using a BioRad Gel Doc EZ imager.

**RNase T2 (Ribonuclease T2)** activity was measured using ssRNA as the substrate. Each reaction mixture (total volume 10 μL) included 1 μg of RNA substrate and 2.5 U RNase T2. The reactions were incubated at 37°C for 15 minutes, then stopped by adding 1 μL of 1% SDS. The reaction products (5 μL) were analyzed by electrophoresis on a 1% agarose gel in 1×TBE buffer, stained with Super GelRed^TM^, run at 120V for 30 minutes, and imaged using a BioRad Gel Doc EZ imager.

**RNase U2 (Ribonuclease U2)** activity was measured using ssRNA as the substrate. Each reaction mixture (total volume 10 μL) included 1× reaction buffer (20 mM sodium acetate, pH 5.0), 1 μg of RNA substrate and 20 U RNase U2. The reactions were incubated at 37°C for 2 hours, then stopped by heating at 75°C for 10 minutes. The reaction products (5 μL) were analyzed by electrophoresis on a 1% agarose gel in 1×TBE buffer, stained with Super GelRed^TM^, run at 120V for 30 minutes, and imaged using a BioRad Gel Doc EZ imager.

**Barnase** activity assay was measured using ssRNA as the substrate. Each reaction mixture (total volume 10 μL) included 1× reaction buffer (30 mM sodium acetate, pH 5.0), 1μg of RNA substrate and free Barnase or Barnase/Barstar complex. Barnase/Barstar were freshly pre-complexed by mixing in equal molar ratio, followed by incubation at room temperature for 10 minutes. The Barnase reactions were incubated at 50°C for 30 minutes, then stopped by adding 500 ng Barstar. The reaction products (5 μL) were analyzed by electrophoresis on a 1% agarose gel in 1×TBE buffer, stained with Super GelRed^TM^, run at 120V for 30 minutes, and imaged using a BioRad Gel Doc EZ imager.

**Human plasma sample collection.** ∼20mL of blood from healthy adult volunteers was drawn with a 23-G needle, immediately transferred to K_3_EDTA collection tubes, and centrifuged for 20 minutes at 2000 g at 4°C. Clear plasma with no observable haemolysis was aliquoted and supplemented with [B_12_H_12_]^2-^ for experiments. Excess plasma was also stored at −80°C if not immediately used.

**Human whole blood sample collection.** ∼20mL of blood from 2 healthy adult volunteers was drawn with a 23-G needle and immediately transferred to K_3_EDTA collection tubes. They were used immediately.

**Human urine sample collection.** ∼50ml of midstream urine was obtained from healthy adult volunteers using a clean-catch technique and used immediately.

### Nucleic acid spike-in assays and (RT)-qPCR

Nucleic acid spike-in was performed by adding λ DNA or MS2 single-stranded RNA to plasma or urine aliquots, with or without nuclease inhibitors. Spiked-in plasma samples were kept at room temperature for various time points. Real-time PCR was performed using TB Green® Premix Ex Taq™ (Tli RNaseH Plus) for DNA spiked-in samples, or One-Step TB Green® PrimeScript™ RT-PCR Kit II (Perfect Real Time) for RNA spiked-in samples, according to the manufacturer’s instructions. The PCR cycle was programmed as follows: (1) 95°C, 1 minute; (2) 95°C, 5 seconds, 60°C, 30 seconds, for 45 cycles; (3) followed by a melting curve analysis at a 0.1°C/second ramping rate ranging from 50°C to 95°C. A 42°C incubation period of 30 minutes was added before PCR initiation for one-step reverse transcription. All biological fluid samples were diluted 100-fold in nuclease-free water to avoid reaction inhibition. All reactions were detected, and Ct values were obtained using the QuantStudio™ 7 Pro Real-Time PCR System (Thermo Fisher Scientific). The primer specificity has been verified using NCBI BLAST to avoid non-specific amplification. Information on primers used is listed in Supp **Table 3**.

### Nucleic acid spike-in and extraction assays

The assays were performed by adding dsDNA PCR product (461 bp), λ DNA, or MS2 single-stranded RNA genome into plasma aliquots, with or without [B_12_H_12_]^2-^. Spiked-in samples were extracted using the standard phenol-chloroform method or the QIAamp ccfDNA/RNA Kit, according to the manufacturer’s instructions. The extracted nucleic acids were quantified by absorbance at 280nm to determine the total amount extracted and the extraction efficiency.

### Human whole blood sample preparation for ccfRNA sequencing

Whole-blood samples from two healthy adult volunteers were stored at room temperature for 0 and 7 days in the presence of various preservatives. At the designated time points, the blood samples were centrifuged for 20 minutes at 2000 g and 4°C to collect the plasma supernatant.

### Haemolysis inhibition assay

The assay was performed by adding a mixture of [B_12_H_12_]^2-^ and zwitterionic compounds to freshly obtained human blood and then incubating at room temperature for 24 hours or 7 days for HEPPSO only. The blood samples were then centrifuged for 10 minutes at 2000 g. The supernatant plasma absorbance was measured at 540nm to quantify the amount of leaked haemoglobin. Plasma supernatant was subject to Nanodrop absorbance measurement at 540nm^[95]^. The residual blood cells were observed using a cell culture microscope at 10x magnification.

### Human plasma ccfRNA sequencing and analysis

Clear plasma samples from three healthy adult volunteers were kept at room temperature for 0 and 7 days in the presence of [B_12_H_12_]^2-^. At the designated time points, plasma nucleic acids were extracted using the QIAamp ccfDNA/RNA Kit according to the manufacturer’s instructions. All extracted RNA samples were deep-frozen in the presence of murine RNase inhibitor during storage and transportation for NGS. Bulk RNA sequencing was outsourced to the BGI Genomics NGS service. RNA quality was examined by capillary electrophoresis in an Agilent 2100 Bioanalyzer. RNA Sequences were aligned with reference to the human genome build NCBI_GCF_000001405.39_GRCh38.p13, from which clean transcript reads were obtained. For each sequencing experiment, the Transcripts Per Million (TPM) was first computed. Transcripts with zero TPM at day 0 were then excluded for each of the control and Na_2_B_12_H_12_ groups. Finally, Kendall’s tau was computed by comparing the results at the day 0 and day 7 time points within each group.

### Nuclear magnetic resonance spectroscopy

NMR titration of barnase in the presence of varying concentrations of Na_2_B_12_H_12_ was performed as previously described ^[76]^. Due to the strong binding of Na_2_B_12_H_12_, an additional six ½ serial dilutions were carried out at and below 50mM Na_2_B_12_H_12_.

### Isothermal titration calorimetry (ITC)

The titration experiments were carried out using a Malvern Panalytical MicroCal PEAQ-ITC. All injections were performed in high-feedback mode at 750 rpm and 25 °C. Sodium dodecahydro-closo-dodecaborate (Na_2_B_12_H_12_) and 2-hydroxypropyl-γ-c-cyclodextrin (2HPγCD) were used throughout the whole titration. 10 mM Na_2_B_12_H_12_ in nuclease-free water was titrated against 1 mM 2HPγCD in nuclease-free water. The initial delay time is 60 s. The initial injection was 0.4 μL, followed by 26 injections of 1.4 μL each, injected every 150 s, for a total of 27 injections. All ITC results were fit to the “One set of sites” model. The stoichiometric number (*N*) and the dissociation constant (*K_d_*) and enthalpy change (*ΔH*) were obtained from Malvern ITC software, and *ΔG* and *TΔS* were calculated from the equation: *ΔG* = – *RT*ln(*Ka*) and *TΔS* = *ΔH* – *ΔG,* respectively. Here, *R* represents the molar gas constant, and *T* denotes the absolute temperature used during the experiment. Raw calorimetric data were analyzed using MicroCal PEAQ-ITC analysis software v1.1.0.1262, and final figures were plotted based on the resulting raw data using GraphPad Prism 10.0.

### Dynamic Viscometry

Viscosity measurements were conducted using a Ubbelohde Viscometer with a 0.24 mm capillary at 20°C. The measured concentrations of boron clusters ranged from 0 M to 0.3125M.

### Electrospray ionization mass spectrometry (ESI-MS)

An AB SCIEX TripleTOF 6600 ESI-MS system was used. 1 mg/mL of Na_2_B_12_H_12_ in nuclease-free water was injected into the mass spectrometer without chromatographic separation, using acetonitrile as the solvent. The injection time was set to 1 minute, and the detection range for the Time-of-Flight (TOF) mass spectrometer was 100-1200 m/z in positive mode.

## Data availability

The data will be made available upon reasonable request to the corresponding author (H.M.L.).

## Supplementary Data

Supplementary Data are available online.

## Acknowledgement

We thank Ms Carmen Chan for administrative assistance in this work.

