## Supplementary Information for "Universal nucleic acid preservation in biological fluids with boron clusters"

### Supplementary Figures

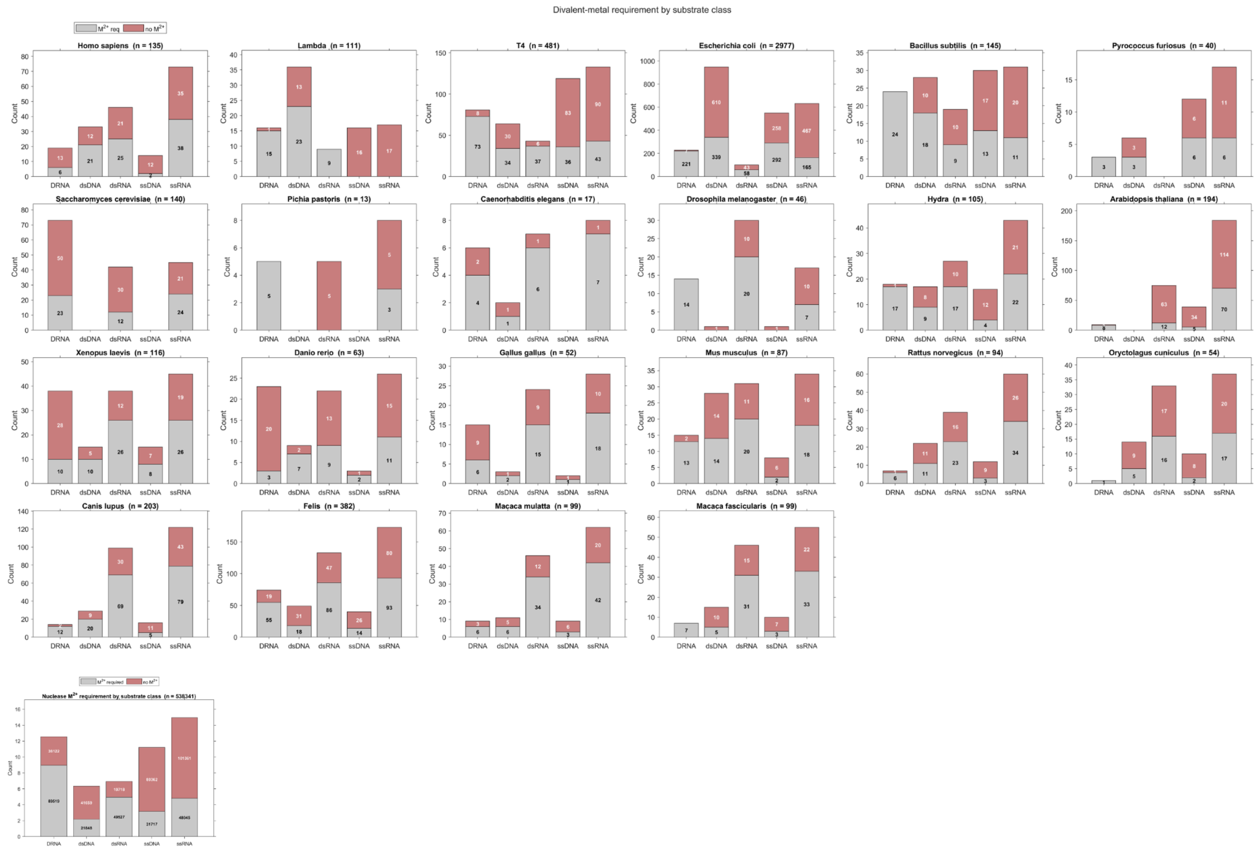


**Supp Fig 1. Divalent ion dependence of all known nucleases in common model organisms.**

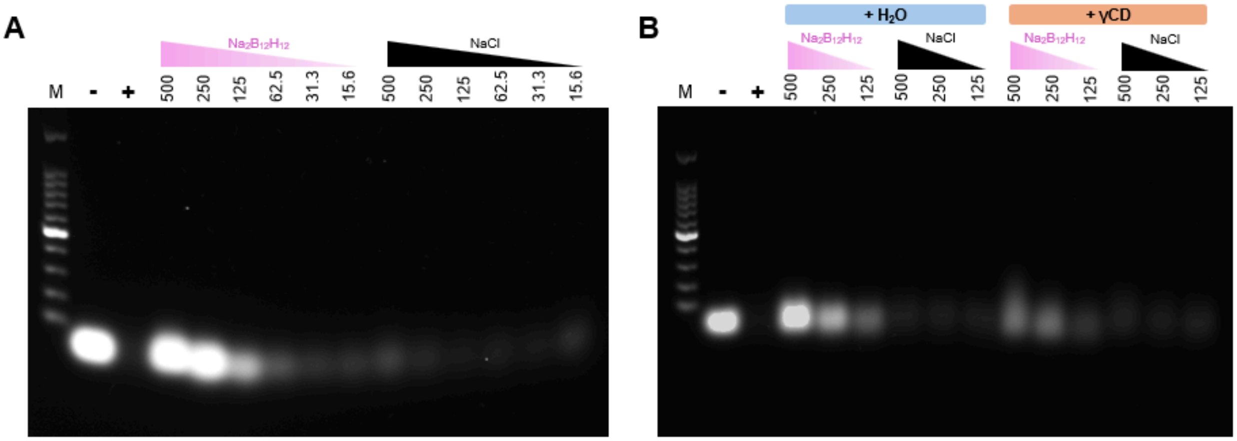


**Supp Fig 2. Inhibition of Exonuclease I by $[B_{12}H_{12}]^{2-}$.** (A) ssDNA was digested by exonuclease I (10 U / 10 μL reaction) in serial dilution of $Na_2B_{12}H_{12}$ or NaCl. (B) $[B_{12}H_{12}]^{2-}$-induced inhibition of nuclease activity was reversed in same molar ratio of γCD. The dilution starting and ending concentrations were as labelled in millimolar. + indicates positive control (substrate digested in recommended condition with no additional additives), - indicates negative control (nuclease-free), and M indicates DNA ladder marker.

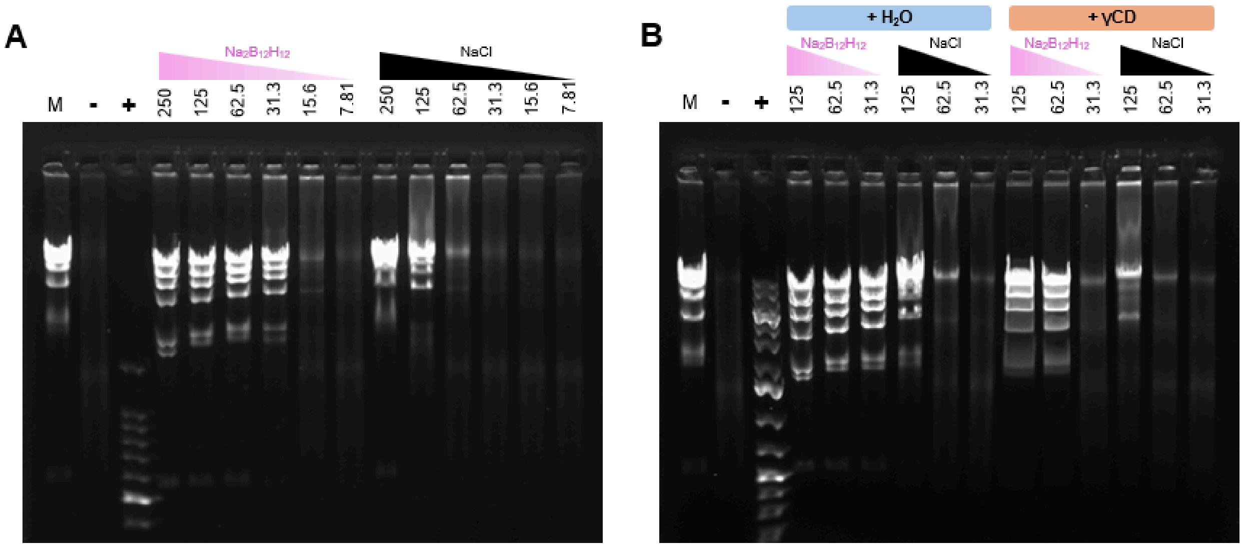


**Supp Fig 3**. **Inhibition of λ exonuclease by $[B_{12}H_{12}]^{2-}$. (A)** λ DNA-HindIII Digest was digested by λ exonuclease (2.5 U / 10 μL reaction) in serial dilution of $Na_2B_{12}H_{12}$ or NaCl. **(B)** $[B_{12}H_{12}]^{2-}$-induced inhibition of nuclease activity was reversed in same molar ratio of γCD. The dilution starting and ending concentrations were as labelled in millimolar. + indicates positive control (substrate digested in recommended condition with no additional additives), - indicates negative control (nuclease-free), and M indicates DNA ladder marker.

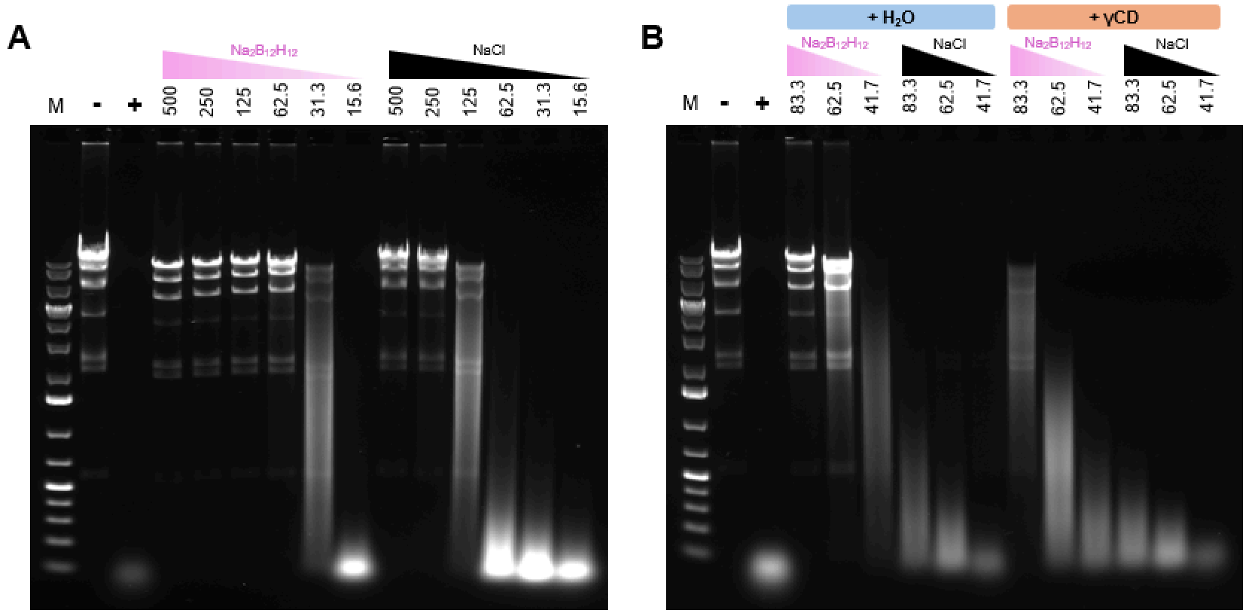


**Supp Fig 4**. **Inhibition of DNase I by $[B_{12}H_{12}]^{2-}$. (A)** λ DNA-HindIII Digest was digested by DNase I (0.5 U / 10 μL reaction) in serial dilution of $Na_2B_{12}H_{12}$ or NaCl. **(B)** $[B_{12}H_{12}]^{2-}$-induced inhibition of nuclease activity was reversed by adding the same molar equivalent of 2HPγCD. The dilution starting and ending concentrations were as labelled in millimolar. + indicates positive control (substrate digested in recommended condition with no additional additives), - indicates negative control (nuclease-free), and M indicates DNA ladder marker.

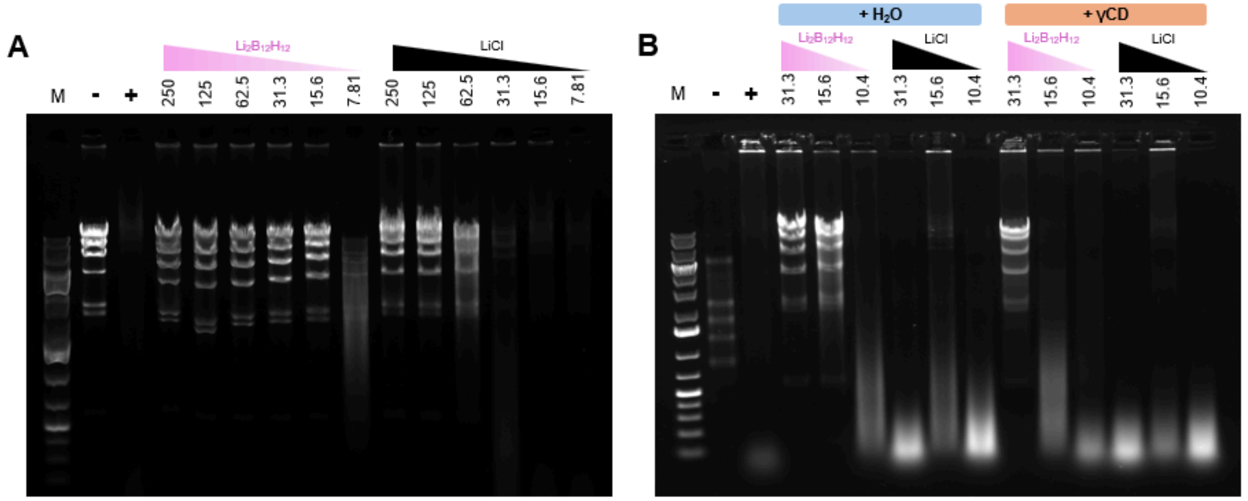


**Supp Fig 5**. **Inhibition of DNase II by $[B_{12}H_{12}]^{2-}$. (A)** λ DNA-HindIII Digest was digested by DNase II (0.5 U / 10 μL reaction) in serial dilution of $Li_2B_{12}H_{12}$ or LiCl. **(B)** $[B_{12}H_{12}]^{2-}$-induced inhibition of nuclease activity was reversed by adding the same molar equivalent of 2HPγCD. The dilution starting and ending concentrations were as labelled in millimolar. + indicates positive control (substrate digested in recommended condition with no additional additives), - indicates negative control (nuclease-free), and M indicates DNA ladder marker.

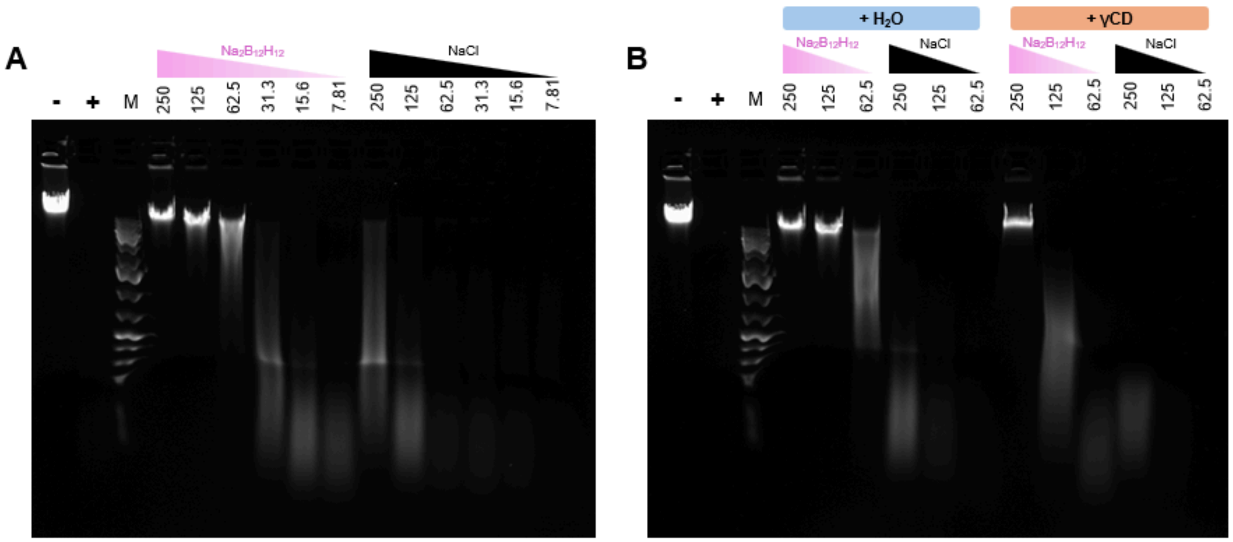


**Supp Fig 6**. **Inhibition of MNase by $[B_{12}H_{12}]^{2-}$. (A)** λ DNA was digested by MNase (1 U / 10 µL reaction) in serial dilution of $Na_2B_{12}H_{12}$ or NaCl. **(B)** $[B_{12}H_{12}]^{2-}$-induced inhibition of nuclease activity was reversed by adding the same molar equivalent of 2HPγCD. The dilution starting and ending concentrations were as labelled in millimolar. + indicates positive control (substrate digested in recommended condition with no additional additives), - indicates negative control (nuclease-free), and M indicates DNA ladder marker.

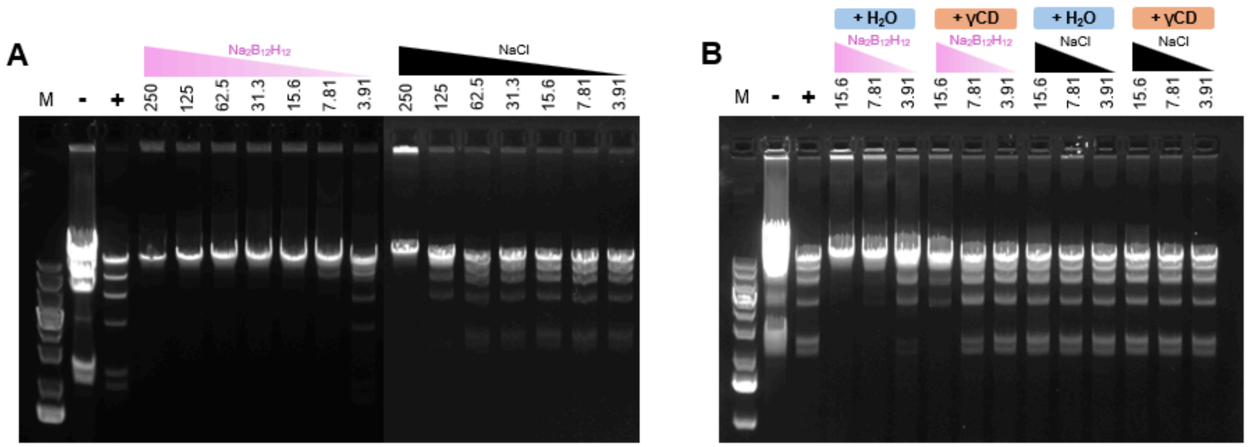


**Supp Fig 7**. **Inhibition of HindIII restriction enzyme by $[B_{12}H_{12}]^{2-}$. (A)** λ DNA was digested by HindIII (10 U / 10 µL reaction) in serial dilution of $Na_2B_{12}H_{12}$ or NaCl. **(B)** $[B_{12}H_{12}]^{2-}$-induced inhibition of nuclease activity was reversed by adding the same molar equivalent of 2HPγCD. The dilution starting and ending concentrations were as labelled in millimolar. + indicates positive control (substrate digested in recommended condition with no additional additives), - indicates negative control (nuclease-free), and M indicates DNA ladder marker.

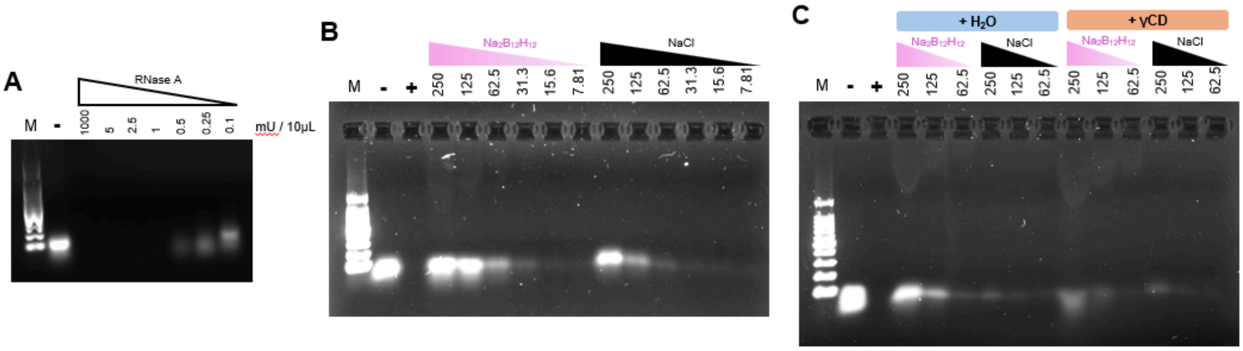


**Supp Fig 8**. **Inhibition of RNase A by $[B_{12}H_{12}]^{2-}$. (A)** RNase A activity titration with ssRNA as the substrate. **(B)** ssRNA was digested by RNase A (50 mU / 10 µL reaction) in serial dilutions of $Na_2B_{12}H_{12}$ or NaCl. **(C)** $[B_{12}H_{12}]^{2-}$-induced inhibition of nuclease activity was reversed by adding the same molar equivalent of 2HPγCD. The dilution starting and ending concentrations were as labelled in millimolar. + indicates positive control (substrate digested in recommended condition with no additional additives), - indicates negative control (nuclease-free), and M indicates DNA ladder marker.

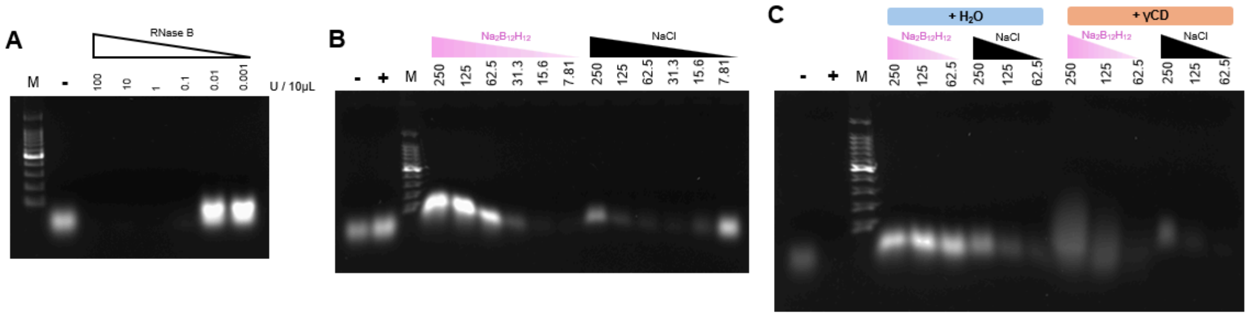


**Supp Fig 9**. **Inhibition of RNase B by $[B_{12}H_{12}]^{2-}$. (A)** RNase B activity titration with ssRNA as the substrate. **(B)** ssRNA was digested by RNase B (0.1 U / 10 µL reaction) in serial dilutions of $Na_2B_{12}H_{12}$ or NaCl. **(C)** $[B_{12}H_{12}]^{2-}$-induced inhibition of nuclease activity was reversed inthe same molar ratio of γCD. The dilution starting and ending concentrations were as labelled in millimolar. + indicates positive control (substrate digested in recommended condition with no additional additives), - indicates negative control (nuclease-free), and M indicates DNA ladder marker.

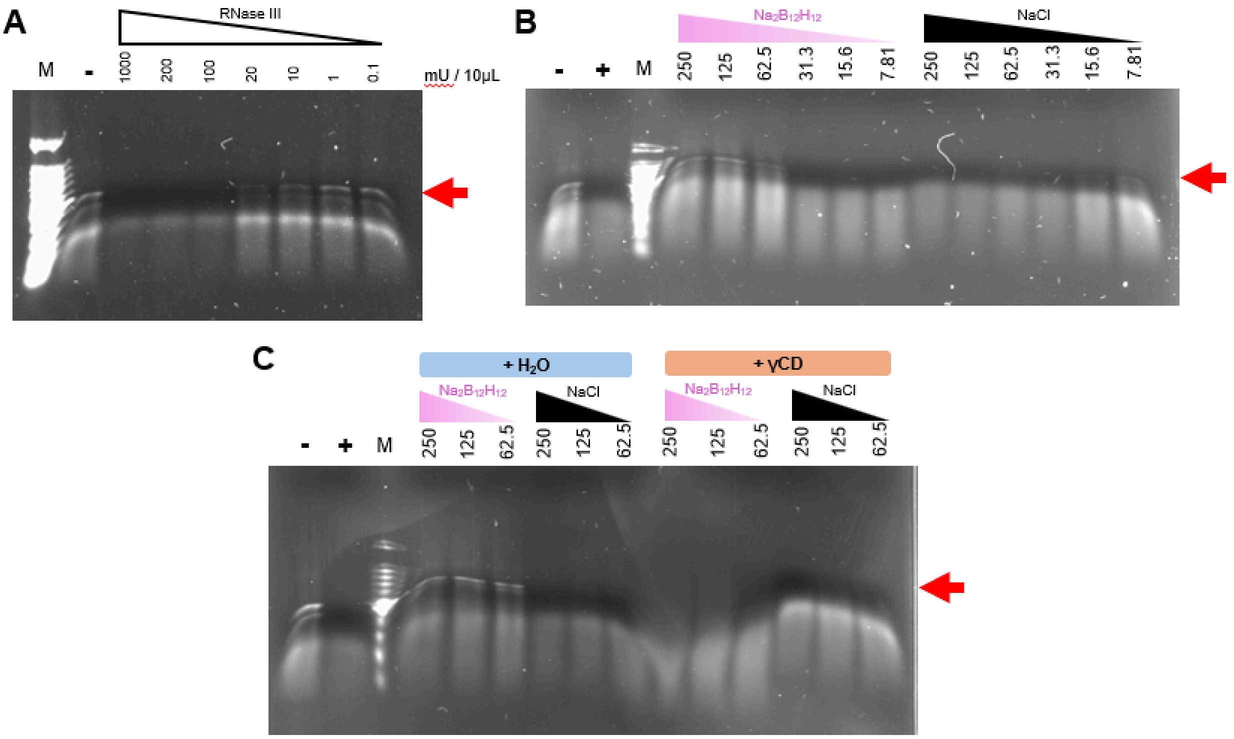


**Supp Fig 10. Inhibition of RNase C (RNase III) by $[B_{12}H_{12}]^{2-}$. (A)** RNase C activity titration with dsRNA ladder as the substrate. **(B)** dsRNA ladder was digested by RNase C (0.1 U / 10 μL reaction) in serial dilutions of $Na_2B_{12}H_{12}$ or NaCl. **(C)** $[B_{12}H_{12}]^{2-}$-induced inhibition of nuclease activity was reversed by adding the same molar equivalent of 2HPγCD. The dilution starting and ending concentrations were as labelled in millimolar. + indicates positive control (substrate digested in recommended condition with no additional additives), - indicates negative control (nuclease-free), and M indicates DNA ladder marker.

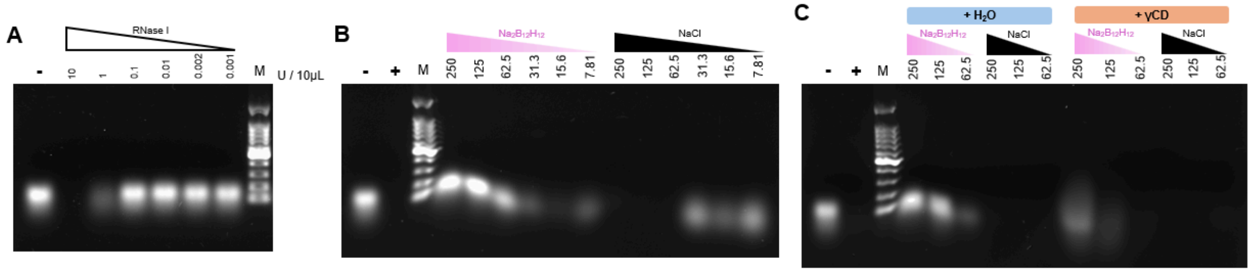


**Supp Fig 11. Inhibition of RNase I by $[B_{12}H_{12}]^{2-}$. (A)** RNase I activity titration with ssRNA as the substrate. **(B)** ssRNA was digested by RNase I (1 U / 10 μL reaction) in serial dilutions of $Na_2B_{12}H_{12}$ or NaCl. **(C)** $[B_{12}H_{12}]^{2-}$-induced inhibition of nuclease activity was reversed by adding the same molar equivalent of 2HPγCD. The dilution starting and ending concentrations were as labelled in millimolar. + indicates positive control (substrate digested in recommended condition with no additional additives), - indicates negative control (nuclease-free), and M indicates DNA ladder marker.

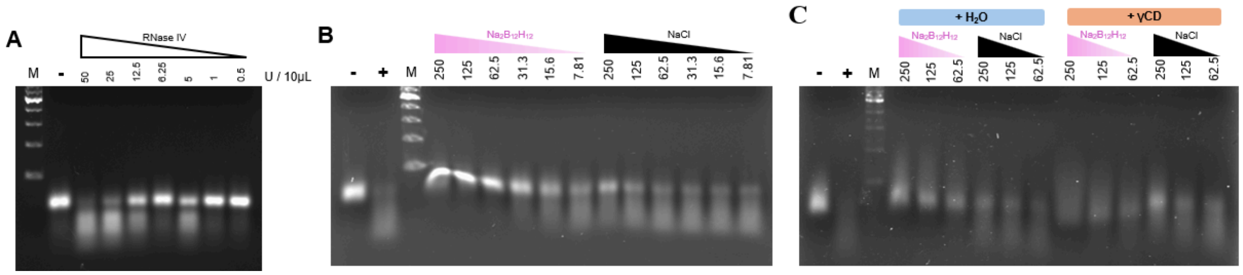


**Supp Fig 12**. **Inhibition of RNase IV by $[B_{12}H_{12}]^{2-}$. (A)** RNase IV activity titration with denatured tRNA as the substrate. **(B)** Denatured tRNA was digested by RNase IV (25 U / 10 µL reaction) in serial dilutions of $Na_2B_{12}H_{12}$ or NaCl. **(C)** $[B_{12}H_{12}]^{2-}$-induced inhibition of nuclease activity was reversed by adding the same molar equivalent of 2HPγCD. The dilution starting and ending concentrations were as labelled in millimolar. + indicates positive control (substrate digested in recommended condition with no additional additives), - indicates negative control (nuclease-free), and M indicates DNA ladder marker.

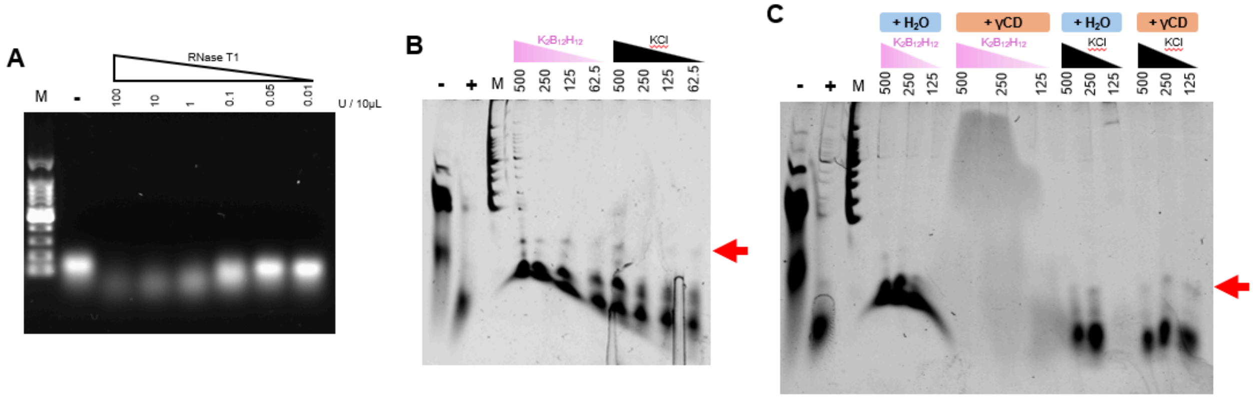


**Supp Fig 13**. **Inhibition of RNase T1 by $[B_{12}H_{12}]^{2-}$. (A)** RNase T1 activity titration with ssRNA as the substrate. **(B)** ssRNA was digested by RNase T1 (1 U / 10 µL reaction) in serial dilutions of $K_2B_{12}H_{12}$ or KCl. **(C)** $[B_{12}H_{12}]^{2-}$-induced inhibition of nuclease activity was reversed by adding the same molar equivalent of 2HPγCD. The dilution starting and ending concentrations were as labelled in millimolar. + indicates positive control (substrate digested in recommended conditions with no additional additives), - indicates negative control (nuclease-free), and M indicates DNA ladder marker.

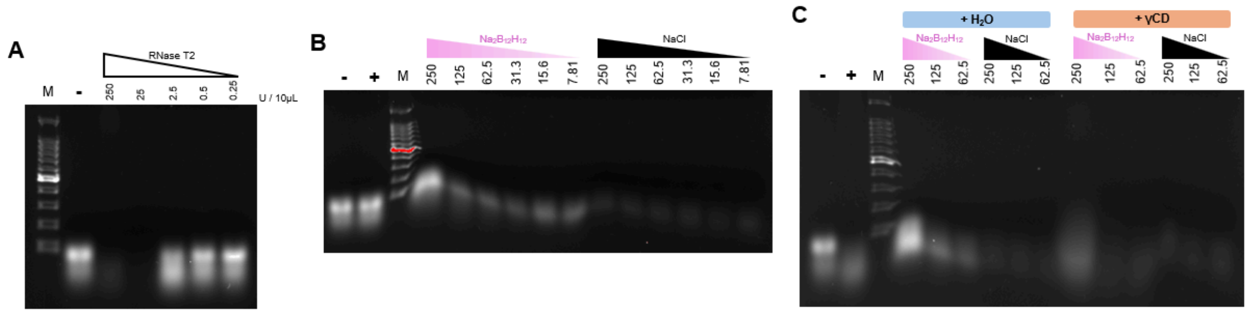


**Supp Fig 14**. **Inhibition of RNase T2 by $[B_{12}H_{12}]^{2-}$. (A)** RNase T2 activity titration with ssRNA as the substrate. **(B)** ssRNA was digested by RNase T2 (2.5 U / 10 µL reaction) in serial dilutions of $Na_2B_{12}H_{12}$ or NaCl. **(C)** $[B_{12}H_{12}]^{2-}$-induced inhibition of nuclease activity was reversed by adding the same molar equivalent of 2HPγCD. The dilution starting and ending concentrations were as labelled in millimolar. + indicates positive control (substrate digested in recommended condition with no additional additives), - indicates negative control (nuclease-free), and M indicates DNA ladder marker.

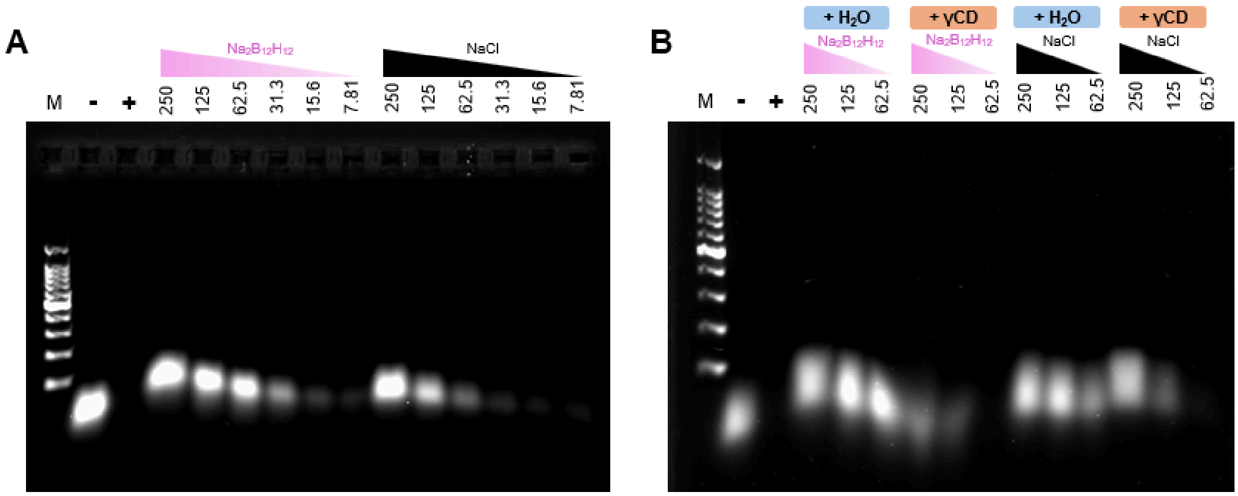


**Supp Fig 15. Inhibition of Nuclease P1 by $[B_{12}H_{12}]^{2-}$. (A)** ssRNA was digested by Nuclease P1 (0.5 U / 10 µL reaction) in serial dilutions of $Na_2B_{12}H_{12}$ or NaCl. **(B)** $[B_{12}H_{12}]^{2-}$-induced inhibition of nuclease activity was reversed by adding the same molar equivalent of 2HPγCD. The dilution starting and ending concentrations were as labelled in millimolar. + indicates positive control (substrate digested in recommended condition with no additional additives), - indicates negative control (nuclease-free), and M indicates DNA ladder marker.

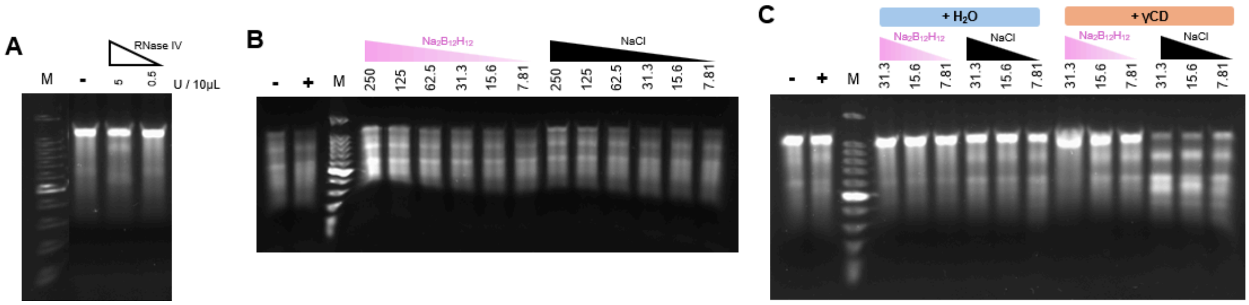


**Supp Fig 16**. **Inhibition of RNase H by $[B_{12}H_{12}]^{2-}$. (A)** RNase H activity titration with RNA-DNA hybrid as the substrate. **(B)** RNA-DNA hybrid was digested by RNase H (5 U / 10 µL reaction) in serial dilutions of $Na_2B_{12}H_{12}$ or NaCl. **(C)** $[B_{12}H_{12}]^{2-}$-induced inhibition of nuclease activity was reversed by adding the same molar equivalent of 2HPγCD. The dilution starting and ending concentrations were as labelled in millimolar. + indicates positive control (substrate digested in recommended condition with no additional additives), - indicates negative control (nuclease-free), and M indicates DNA ladder marker.

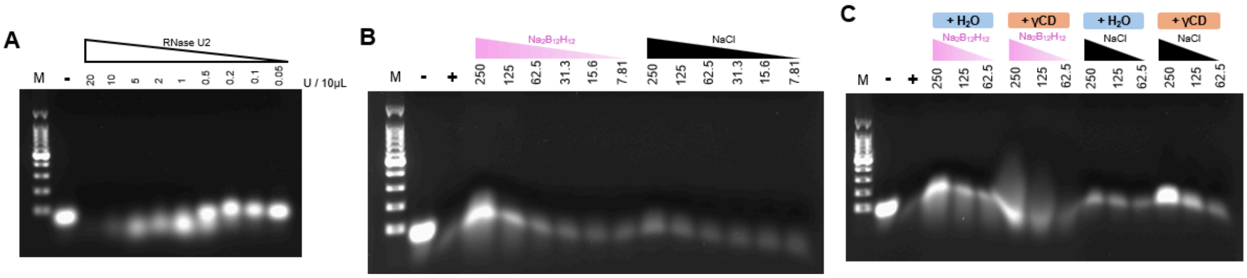


**Supp Fig 17**. **Inhibition of RNase U2 by $[B_{12}H_{12}]^{2-}$. (A)** RNase U2 activity titration with tRNA as the substrate. **(B)** tRNA was digested by RNase U2 (20 U / 10 µL reaction) in serial dilutions of $Na_2B_{12}H_{12}$ or NaCl. **(C)** $[B_{12}H_{12}]^{2-}$-induced inhibition of nuclease activity was reversed by adding the same molar equivalent of 2HPγCD. The dilution starting and ending concentrations were as labelled in millimolar. + indicates positive control (substrate digested in recommended condition with no additional additives), - indicates negative control (nuclease-free), and M indicates DNA ladder marker.

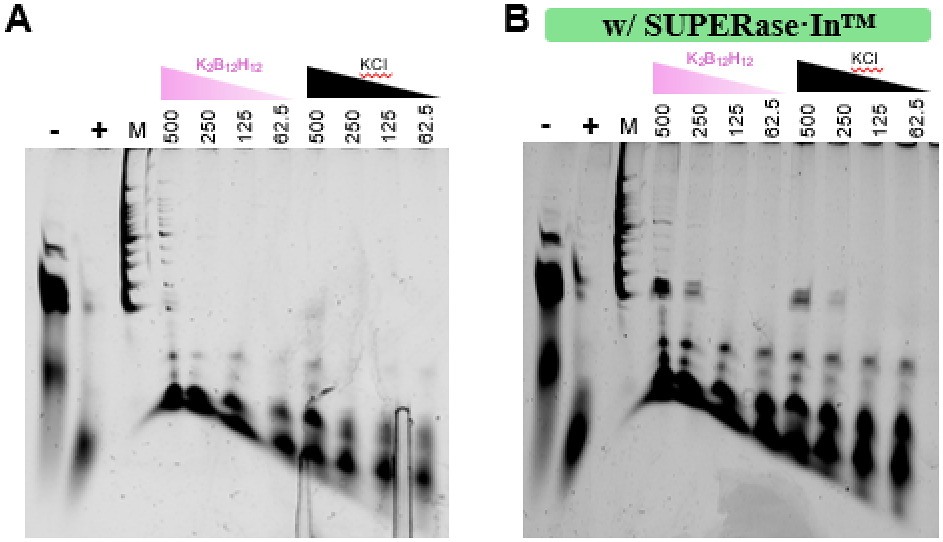


**Supp Fig 18**. **$[B_{12}H_{12}]^{2-}$ provide an additional RNase inhibition effect for RNase T1 on top of SUPERase·In™**. **(A)** ssRNA was digested by RNase T1 (1 U / 10 µL reaction) in serial dilutions of $K_2B_{12}H_{12}$ or KCl. **(B)** The addition of $[B_{12}H_{12}]^{2-}$ further protected RNA from RNase T1 digestion than SUPERase·In™ alone. The dilution starting and ending concentrations were as labelled in millimolar. + indicates positive control (substrate digested in recommended conditions with no additional additives), - indicates negative control (nuclease-free), and M indicates DNA ladder marker.

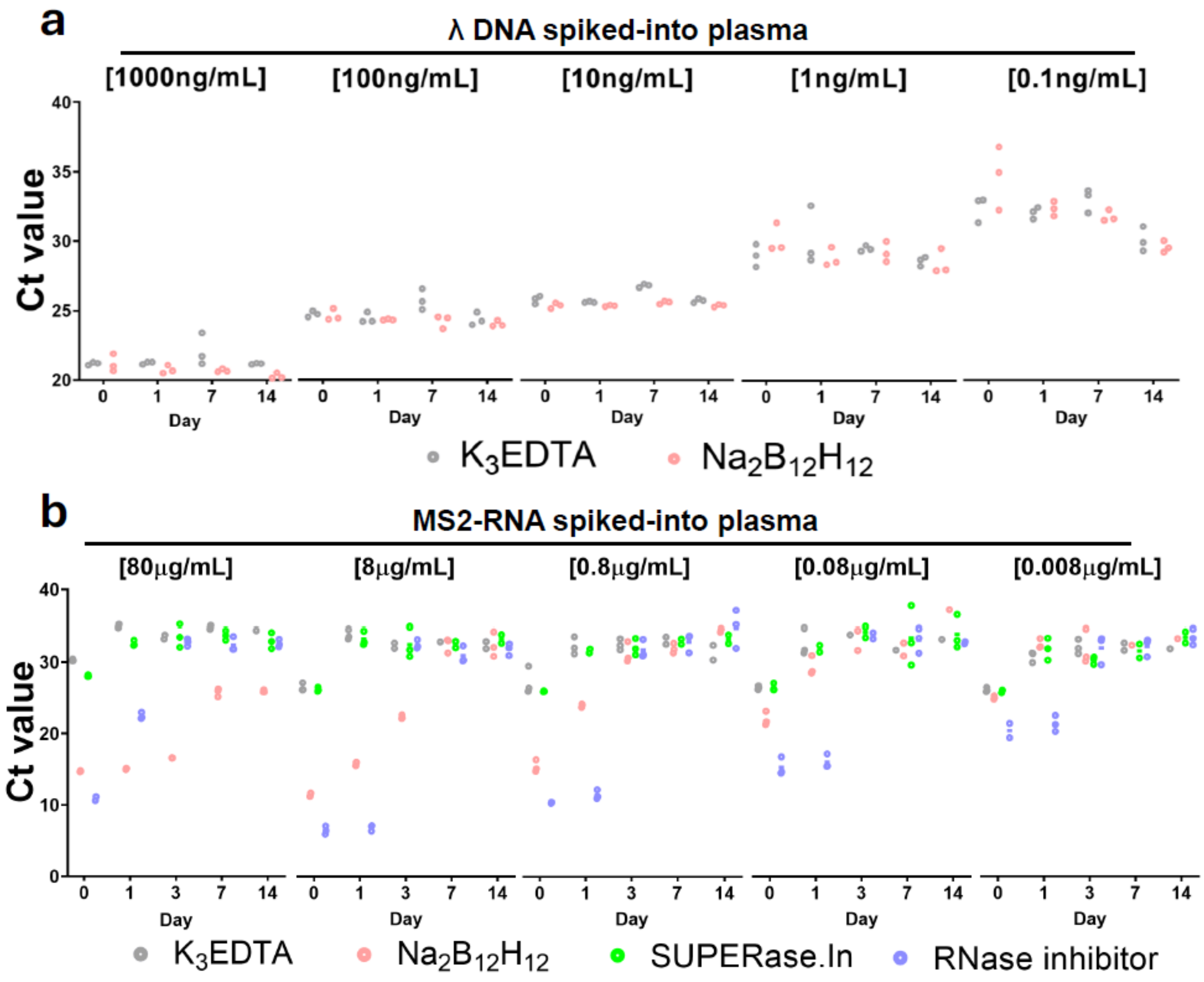


**Supp Fig 19**. **$[B_{12}H_{12}]^{2-}$ preserves λ DNA and MS2 RNA in human plasma at room temperature. a** Plasma λ DNA levels were measured at various time-points (0, 1, 7 and 14 days) by real-time PCR (n=3). Final plasma concentration of $[B_{12}H_{12}]^{2-}$ is 0.25 M, and the initial λ DNA spike-in concentration in plasma is as indicated. A circle represents an individual Ct value. **b** Plasma MS2 ssRNA levels were measured at various time-points (0, 1, 3, 7 and 14 days) by real-time PCR (n=3). Final plasma concentrations of $[B_{12}H_{12}]^{2-}$, SUPERase.In and RNase inhibitor are 0.25 M, 2U/μL and 4U/μL, respectively. The initial MS2 ssRNA spike-in concentration in plasma is as indicated. A circle represents an individual Ct value.

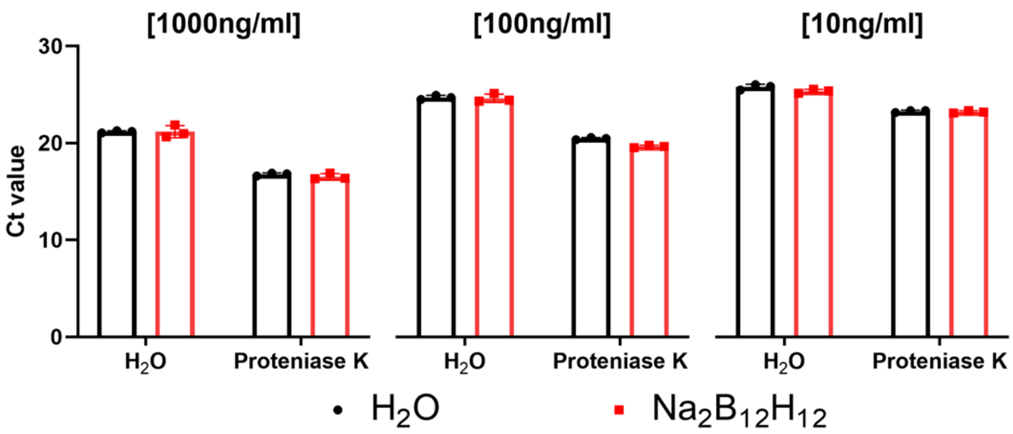


**Supp Fig 20**. **Proteinase K digestion of human plasma leads to a reduction in Ct value for spiked-in DNA**. Plasma λ DNA levels were measured after Proteinase K digestion (0.8U / rxn, 37 °C for 30 min followed by 95 °C inactivation for 10 min) by real-time PCR (n=3). Final plasma concentration of $[B_{12}H_{12}]^{2-}$ is 0.25 M, and the initial λ DNA spike-in concentration in plasma is as indicated above each graph in square brackets.

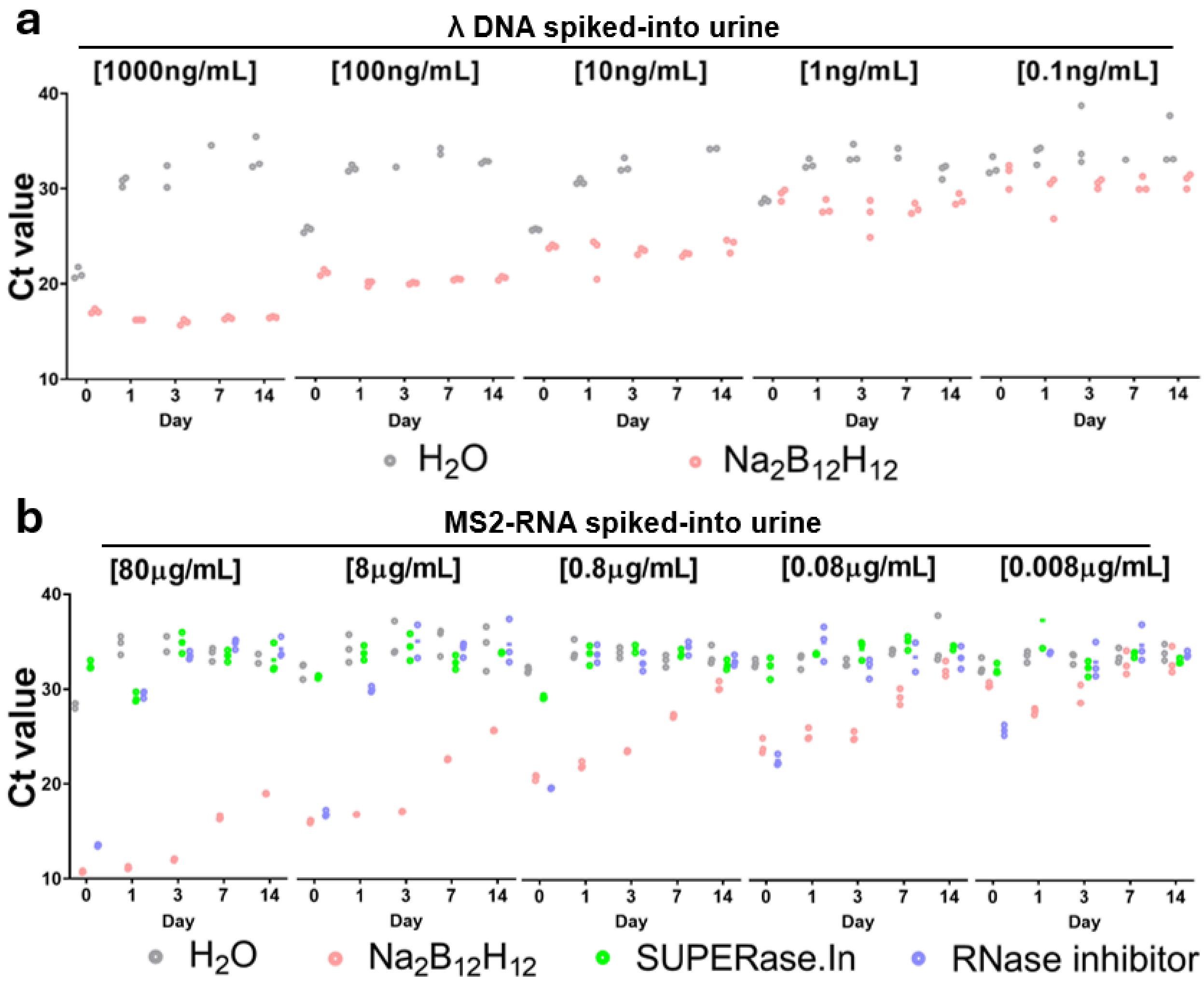


**Supp Fig 21**. **$[B_{12}H_{12}]^{2-}$ preserves λ DNA and MS2 RNA in human urine at room temperature. a** Plasma λ DNA levels were measured at various time-points (0, 1, 3, 7 and 14 days) by real-time PCR (n=3). Final plasma concentration of $[B_{12}H_{12}]^{2-}$ is 0.25 M, and the initial λ DNA spike-in concentration in urine is as indicated. A circle represents an individual Ct value. **b** Plasma MS2 ssRNA levels were measured at various time-points (0, 1, 3, 7 and 14 days) by real-time PCR (n=3). Final plasma concentrations of $[B_{12}H_{12}]^{2-}$, SUPERase.In and RNase inhibitor are 0.25 M, 2U/μL and 4U/μL, respectively. The initial MS2 ssRNA spike-in concentration in plasma is as indicated. A circle represents an individual Ct value.

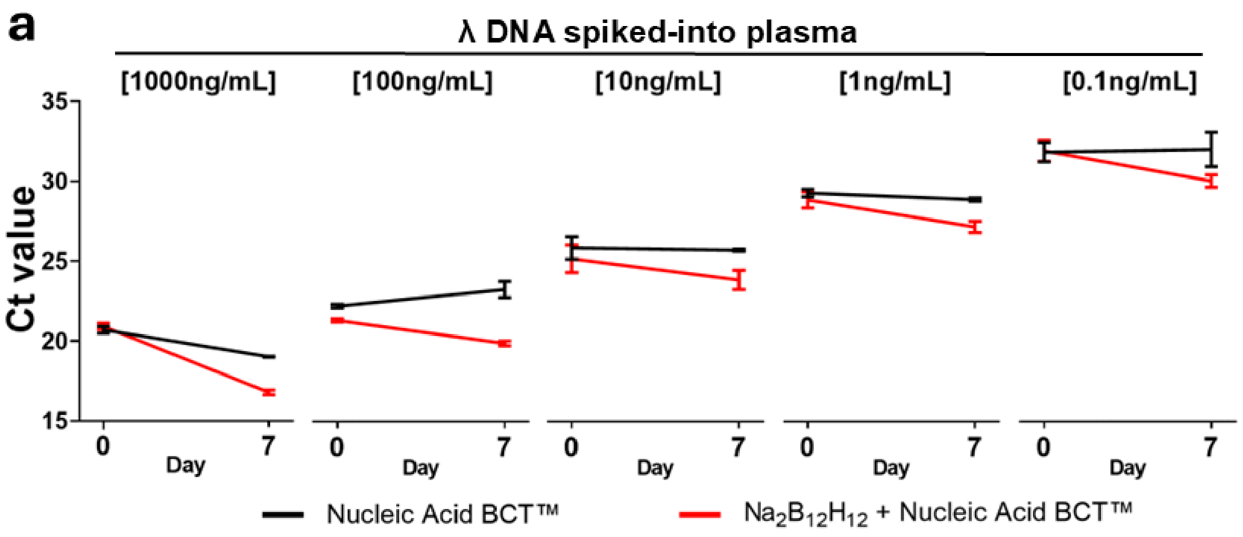


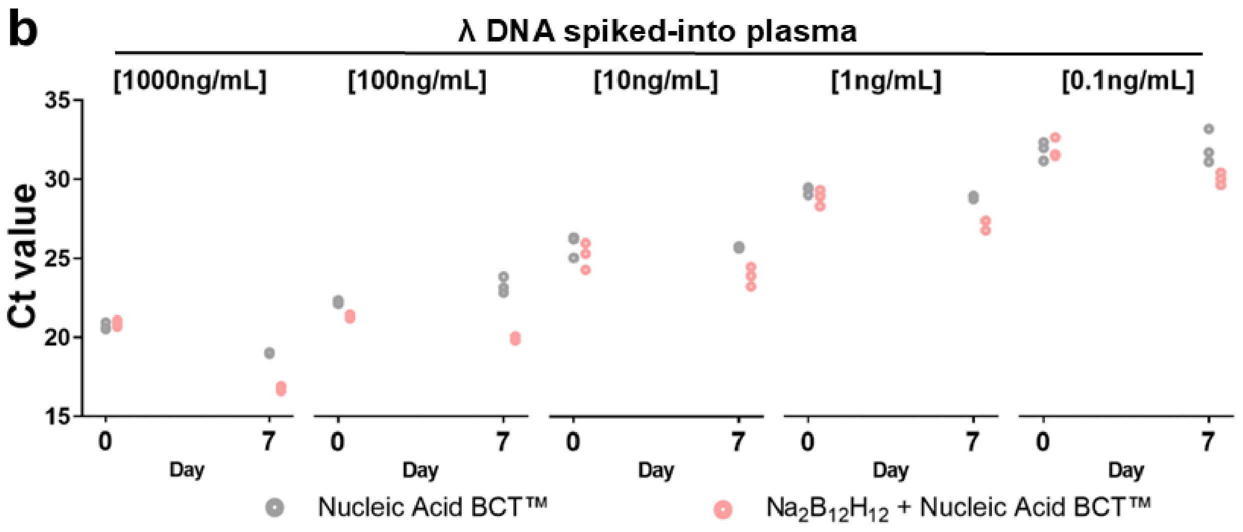


**Supp Fig 22. Comparison between $[B_{12}H_{12}]^{2-}$ and Streck Nucleic Acid BCT$^{TM}$ on the effect of λ DNA preservation in human plasma at room temperature.** Plasma λ DNA levels were measured on days 0 and 7 by real-time PCR (n=3). Final plasma concentration of $[B_{12}H_{12}]^{2-}$ and Streck Nucleic Acid BCT$^{TM}$ are 0.25 M and 1 X, respectively. The initial λ DNA spike-in concentration in plasma is as indicated. **a** Bars represent group mean ± SD. **b** Circles represent the individual Ct values of each sample.

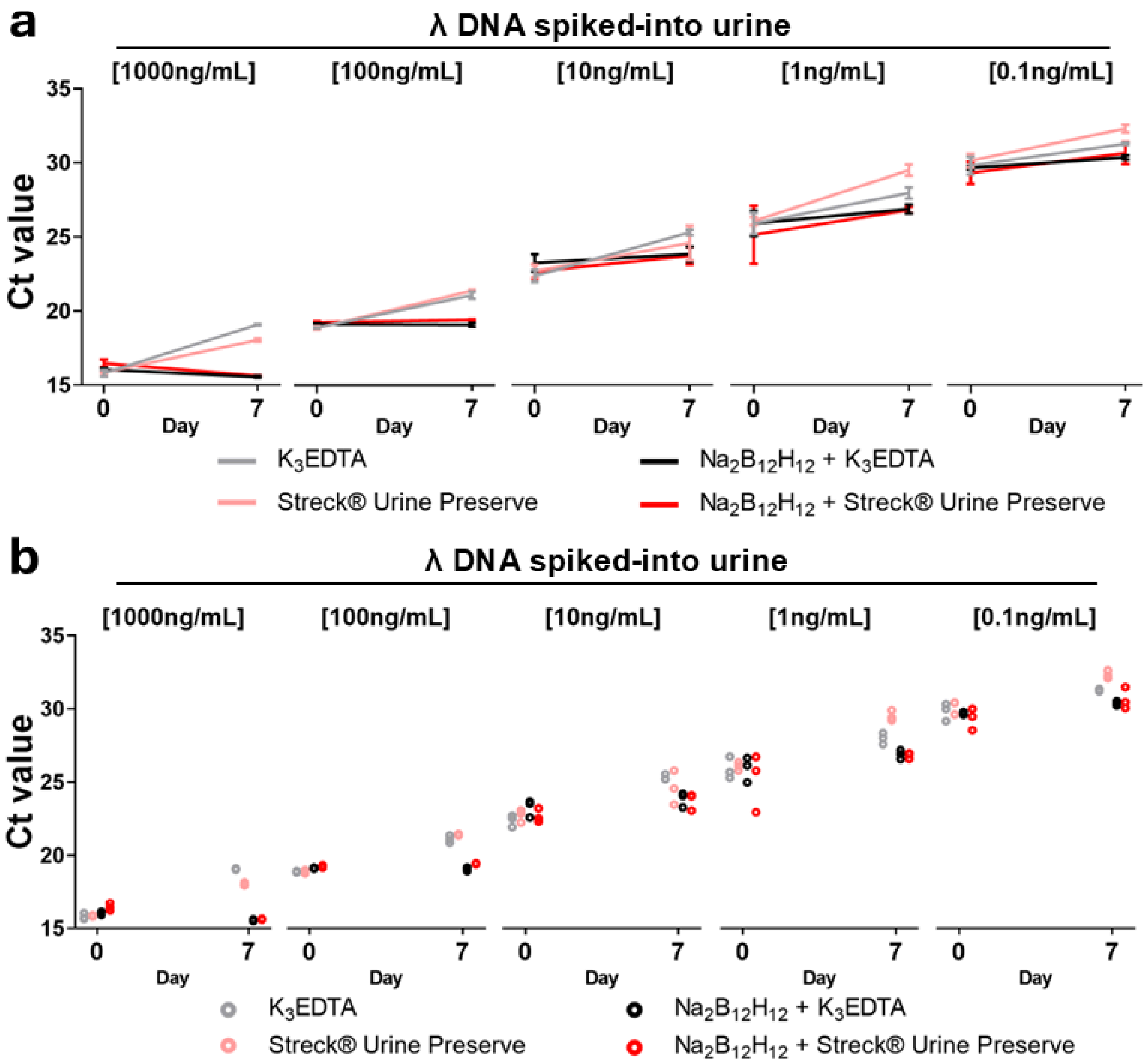


**Supp Fig 23**. **Comparison between $[B_{12}H_{12}]^{2-}$, Streck® Urine Preserve and $K_3EDTA$ on the effect of λ DNA preservation in human urine at room temperature.** Urine λ DNA levels were measured on days 0 and 7 by real-time PCR (n=3). The final urine concentration of $[B_{12}H_{12}]^{2-}$, Streck Urine Preserve, and $K_3EDTA$ is 0.25 M, 1 ×, and 1.5 mg/mL, respectively. The initial λ DNA spike-in concentration in urine is as indicated. **a** Bars represent group mean ± SD. **b** Circles represent the individual Ct values of each sample.

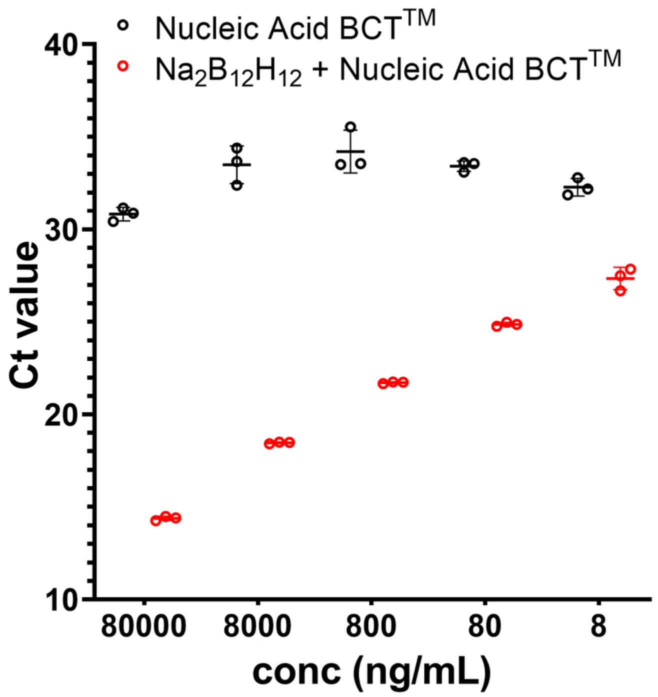


**Supp Fig 24**. **Comparison between $[B_{12}H_{12}]^{2-}$ and Streck Nucleic Acid BCT™ on the effect of MS2 RNA preservation in human plasma at room temperature.** MS2 RNA levels were measured on day 0 by real-time PCR (n=3). Final plasma concentration of $[B_{12}H_{12}]^{2-}$ and Streck Nucleic Acid BCT**™** are 0.25 M and 1 X , respectively. The initial MS2 RNA spike-in concentration in plasma is as indicated. Bars represent group mean ± SD. Circles represent the individual Ct values of each sample.

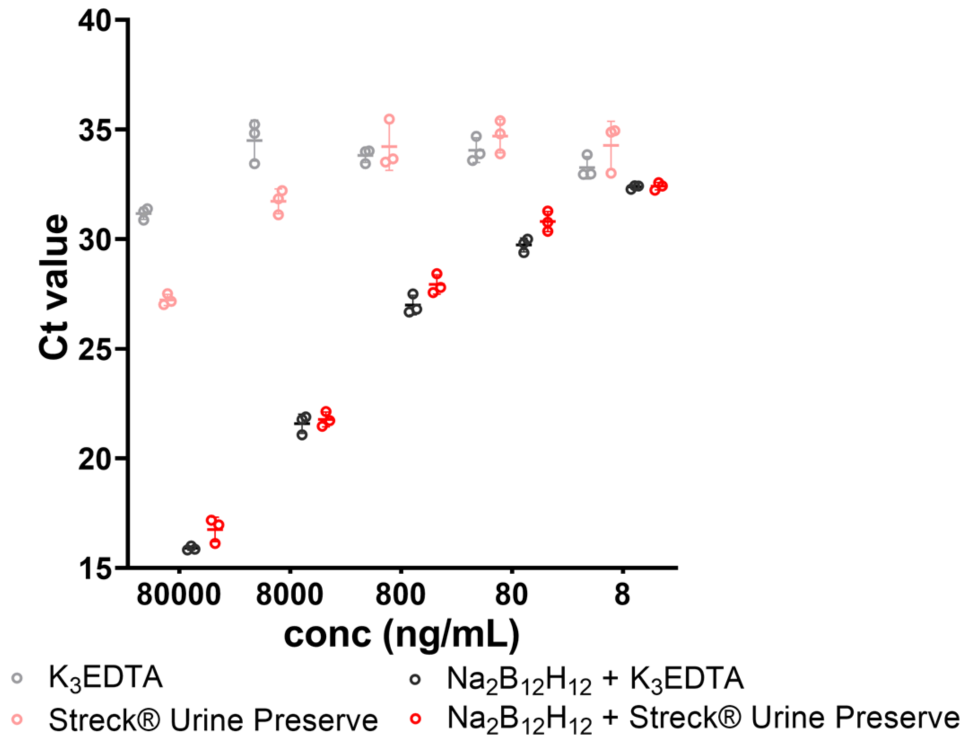


**Supp Fig 25**. **Comparison between $[B_{12}H_{12}]^{2-}$, Streck® Urine Preserve and $K_3$EDTA on the effect of MS2 RNA preservation in human urine at room temperature.** MS2 RNA levels were measured on day 0 by real-time PCR (n=3). Final plasma concentration of $[B_{12}H_{12}]^{2-}$ and Streck Nucleic Acid BCT™ are 0.25 M and 1x, respectively. The initial MS2 RNA spike-in concentration in plasma is as indicated. Bars represent group mean ± SD. Circles represent the individual Ct values of each sample.

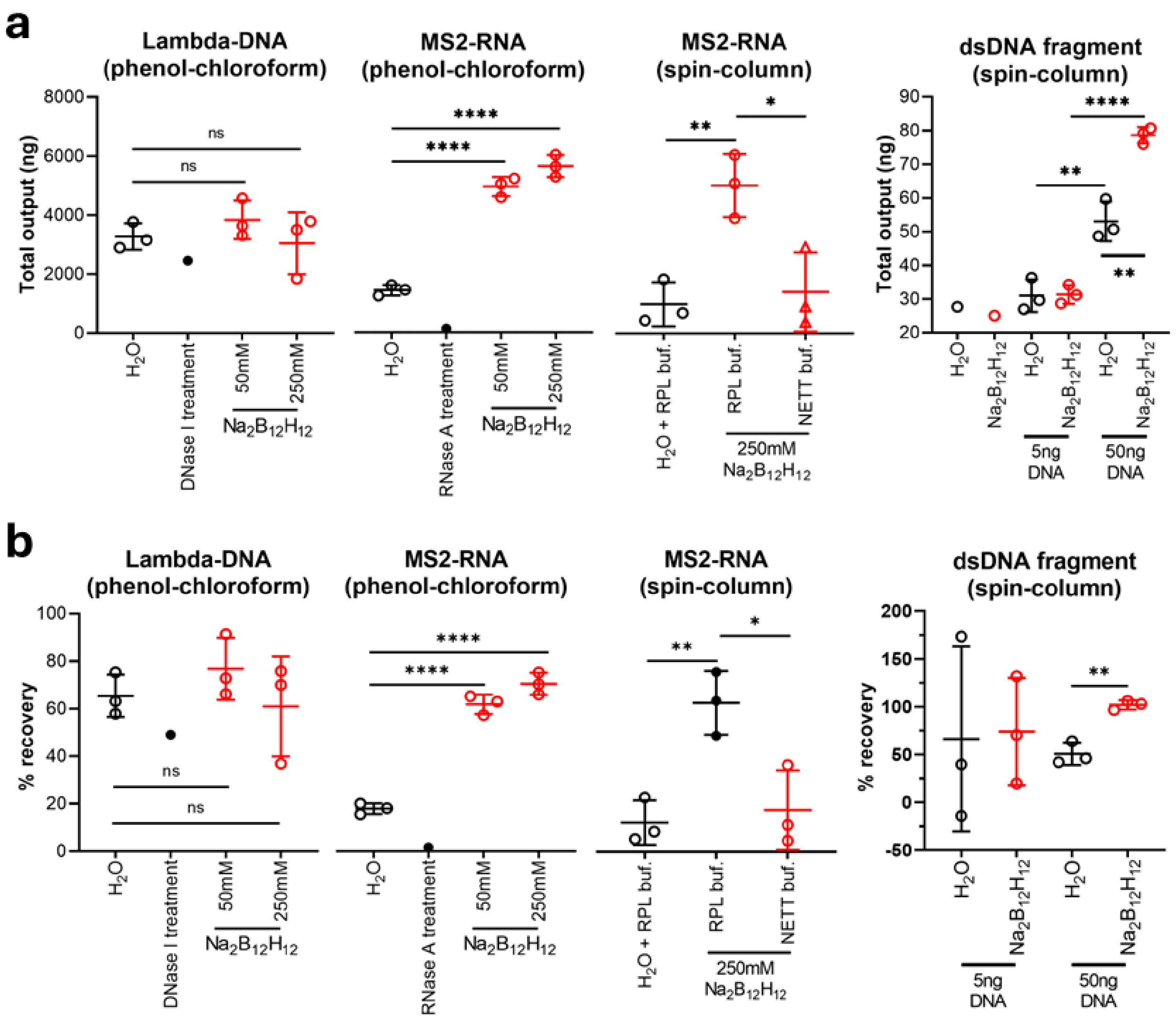


**Supp Fig 26. Effects of $Na_2B_{12}H_{12}$ on the extraction of nucleic acids from human plasma. a** Total amount of nucleotide extracted, and **b** extraction recovery rate, of lambda-DNA, MS2-ssRNA or dsDNA fragment spiked into human plasma. Nucleotide extraction was performed with the phenol-chloroform method or QIAGEN QIAamp ccfDNA/RNA Kit (spin-column). RPL buffer was supplied in the Kit, while NETT buffer (0.1M $Na_2B_{12}H_{12}$, 10mM K3EDTA, 1% Tween-20 and 50mM Tris-HCl, pH 7.5) was homemade. n.s., not significant; *: $P < 0.05$; **: $P < 0.01$; ****: $P < 0.0001$ by unpaired two-tailed Student's *t*-test.

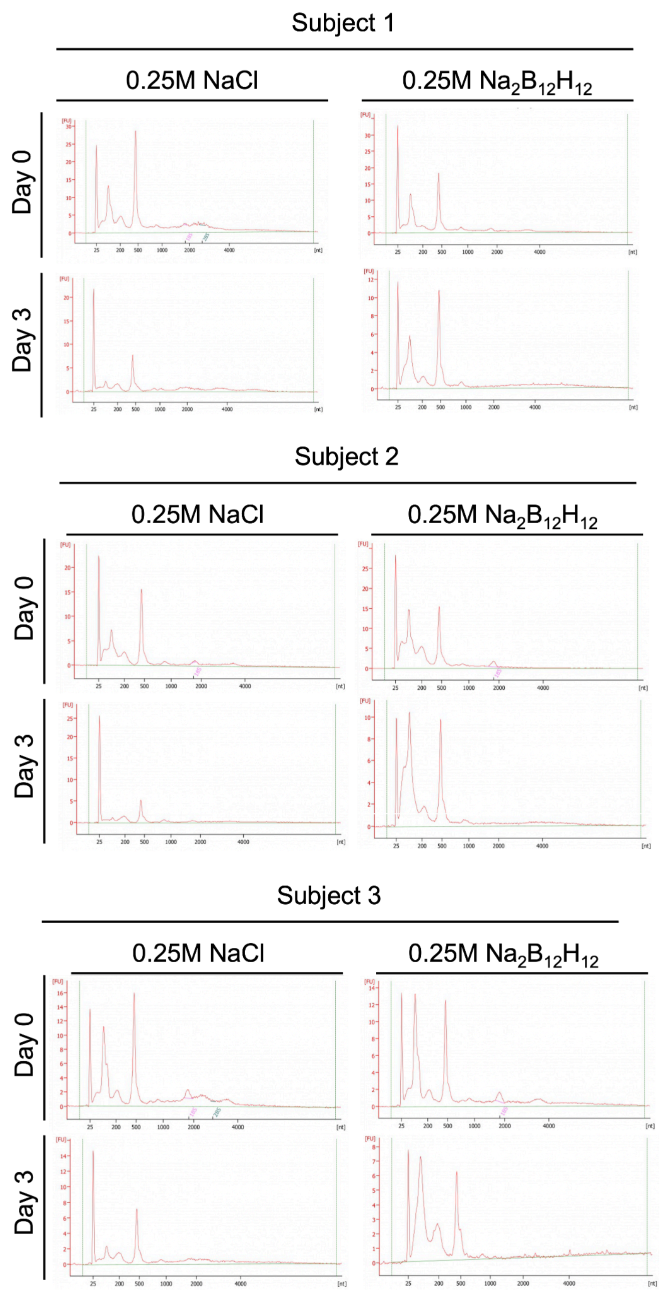


**Supp Fig 27. Capillary electropherogram of plasma nucleic acids upon prolonged storage.** Plasma from 3 subjects were used and incubated in the absence or presence of $Na_2B_{12}H_{12}$ over 7 days at ambient temperatures (15-25℃), all samples were collected using standard $K_3EDTA$ tubes.

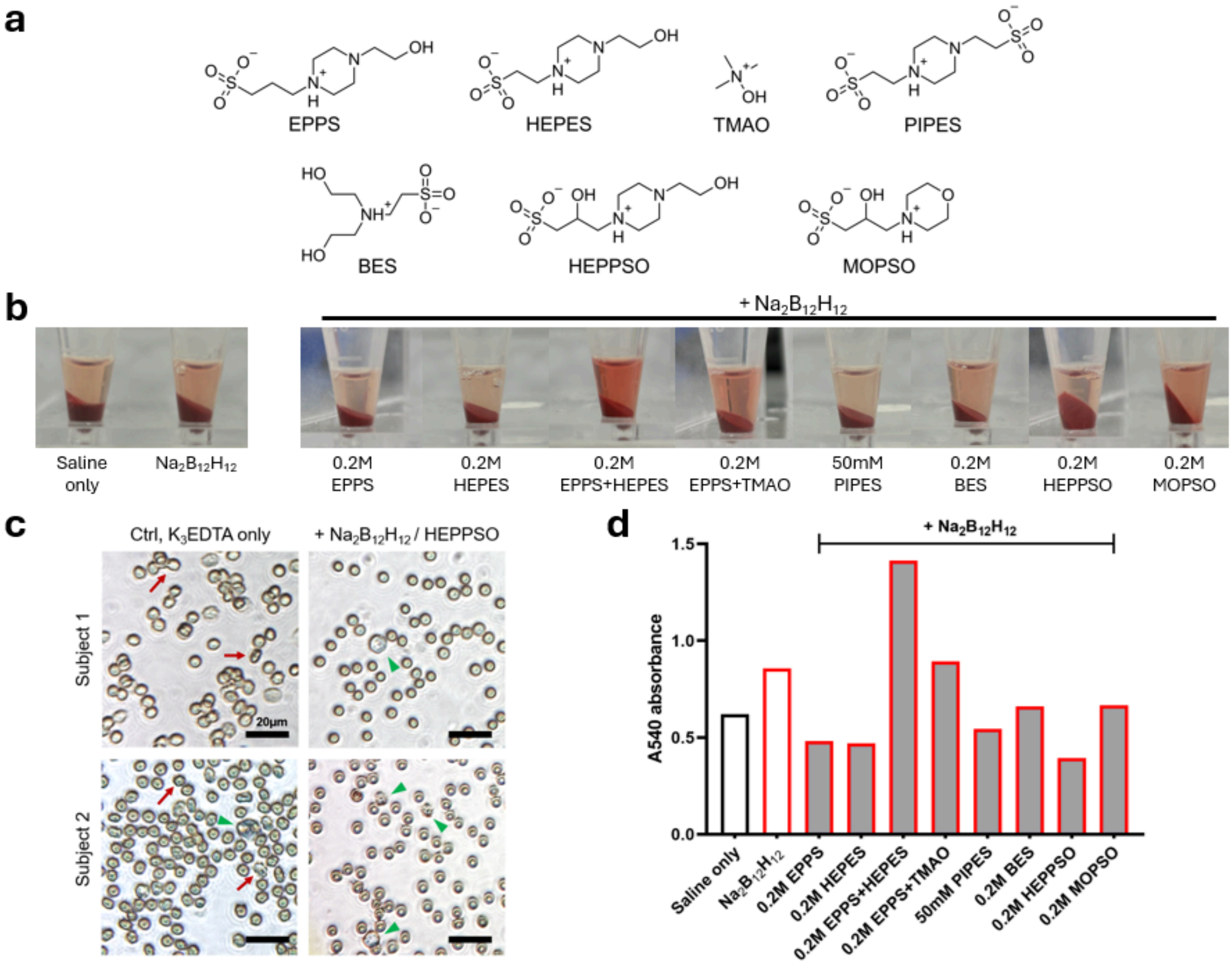


**Supp Fig 28. Screening zwitterionic compounds to prevent $[B_{12}H_{12}]^{2-}$-induced hemolysis. A.** Structures of the zwitterionic compounds tested. EPPS: 3-[4-(2-Hydroxyethyl)piperazin-1-yl]propane-1-sulfonic acid; HEPES: 4-(2-Hydroxyethyl)-1-piperazineethane sulfonic acid; TMAO: Trimethylamine N-oxide; PIPES: 1,4-Piperazine bis(propanesulfonic acid); BES: N,N-Bis(2-hydroxyethyl)-2-aminoethanesulfonic acid; HEPPSO: 4-(2-Hydroxyethyl)piperazine-1-(2-hydroxypropane-3-sulfonic acid); MOPSO: 2-Hydroxy-4-morpholinepropanesulphonic acid. **b.** Photographs of whole blood stored in 50 mM $Na_2B_{12}H_{12}$ in the presence or absence of zwitterionic compounds. **c** Microphotograph of red blood cells (RBC) after 1 day of storage in the presence or absence of $Na_2B_{12}H_{12}$ / HEPPSO. Red arrows indicate crumpled, distorted-looking RBCs. White blood cells (green arrowheads) are seen more frequently in the $Na_2B_{12}H_{12}$/HEPPSO group, which are otherwise known to be sensitive to lysis upon prolonged storage. **d** 540 nm spectrophotometry of the supernatant plasma after 7 days storage of whole blood in the presence of $[B_{12}H_{12}]^{2-}$ and zwitterionic compounds.

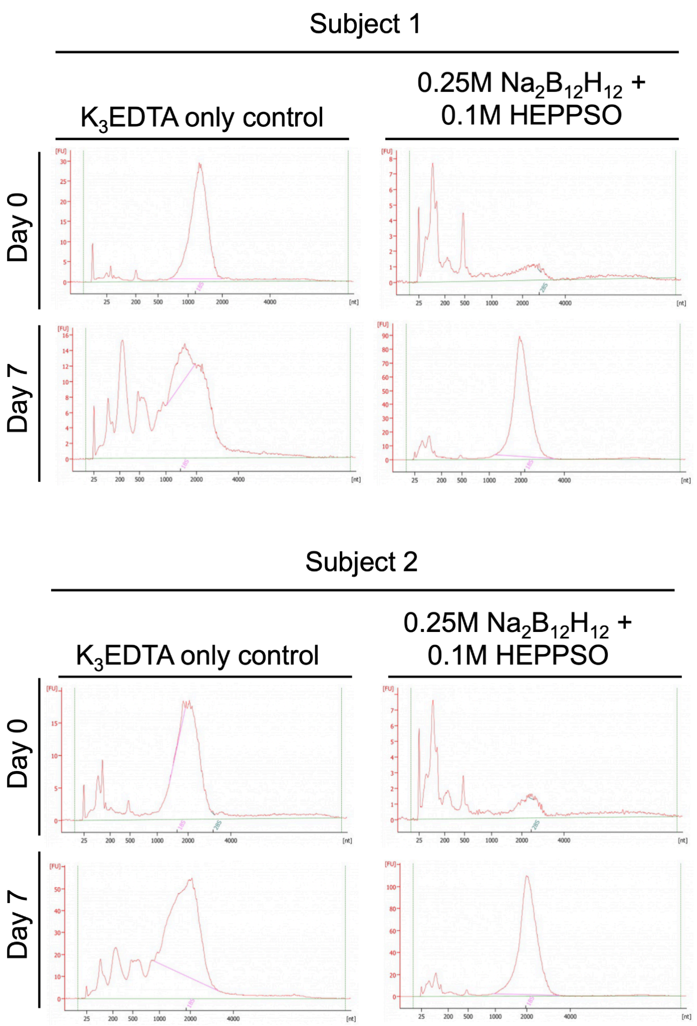


**Supp Fig 29. Capillary electropherograms of extracted nucleic acids upon prolonged storage as whole blood.** Plasma from 2 subjects was incubated in the absence or presence of $Na_2B_{12}H_{12}$/HEPPSO for 7 days at ambient temperature (15-25℃); all samples were collected in standard $K_3EDTA$ tubes.

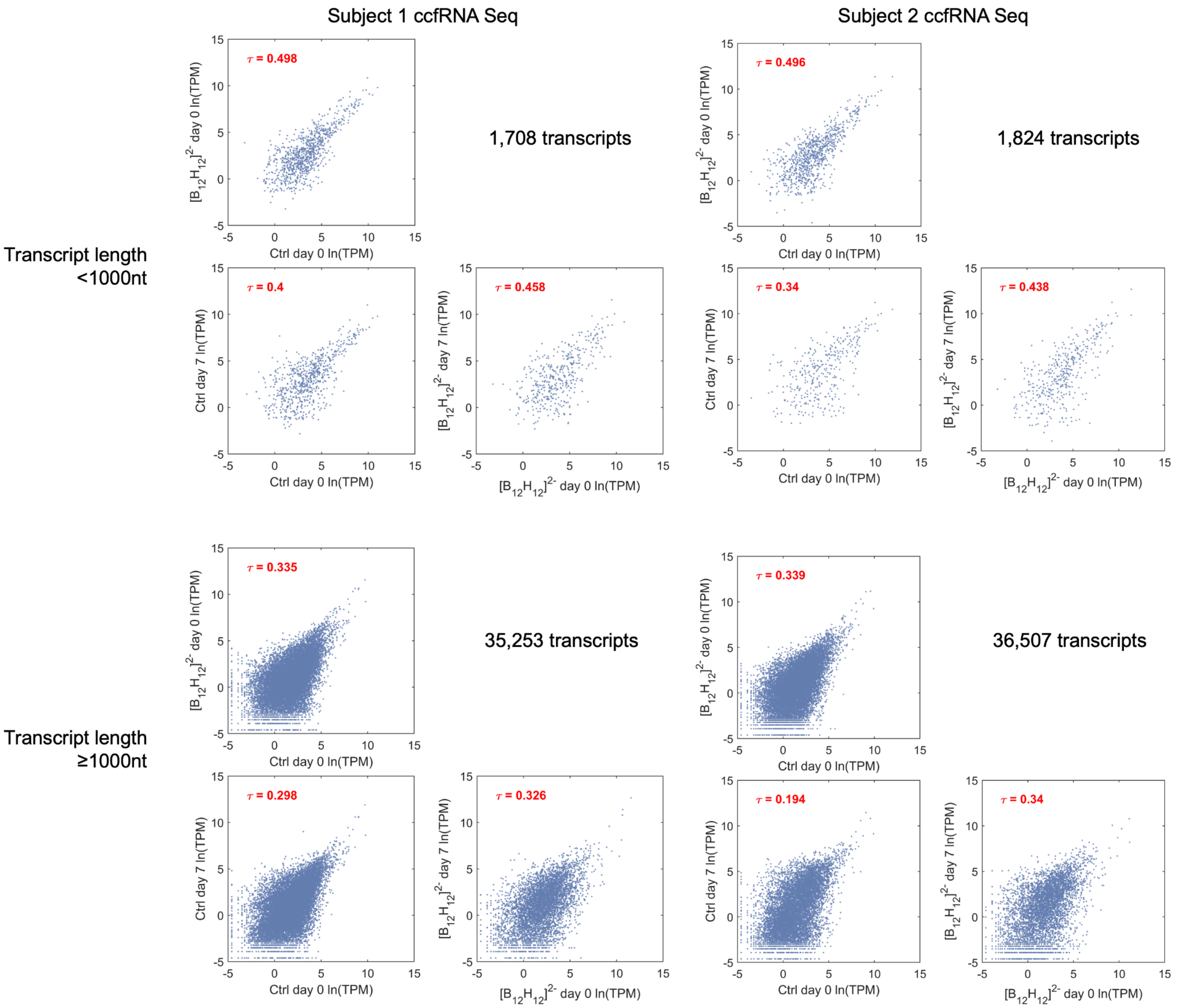


**Supp Fig 30. ccfRNA Sample correlation by sequenced transcript-level TPM over 7 days of storage under different conditions.** The analysis was performed as in **Figure 6e**, but categorised by transcript length using a 1000 nt cut-off.

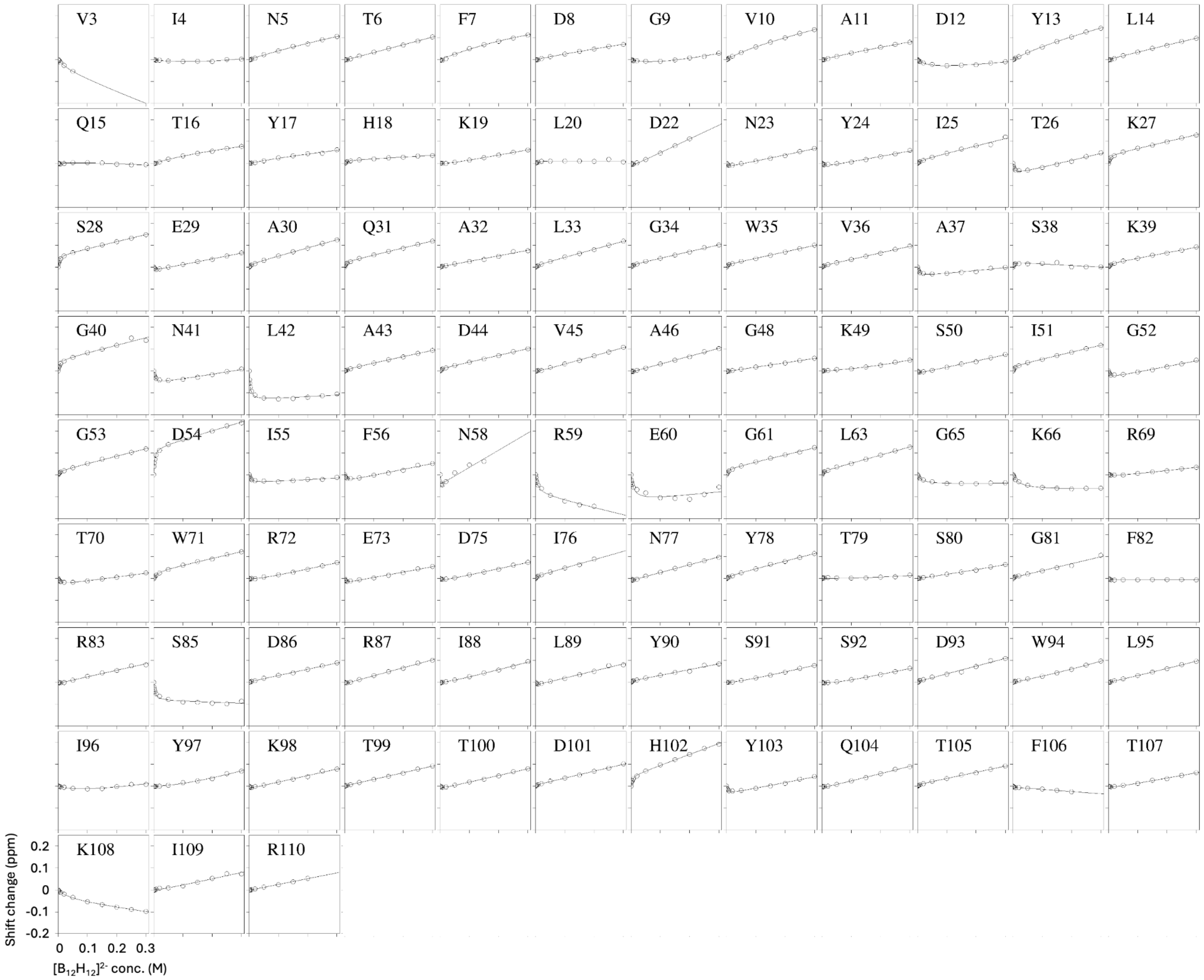


**Supp Fig 31. $^{1}H_{N}$ NMR shifts of barnase residues with varying concentrations of $[B_{12}H_{12}]^{2-}$.** Chemical shift scale and concentration were shown at the bottom left.

$Na_2B_{12}H_{12}$
M.W. = 187.81 g mol$^{-1}$

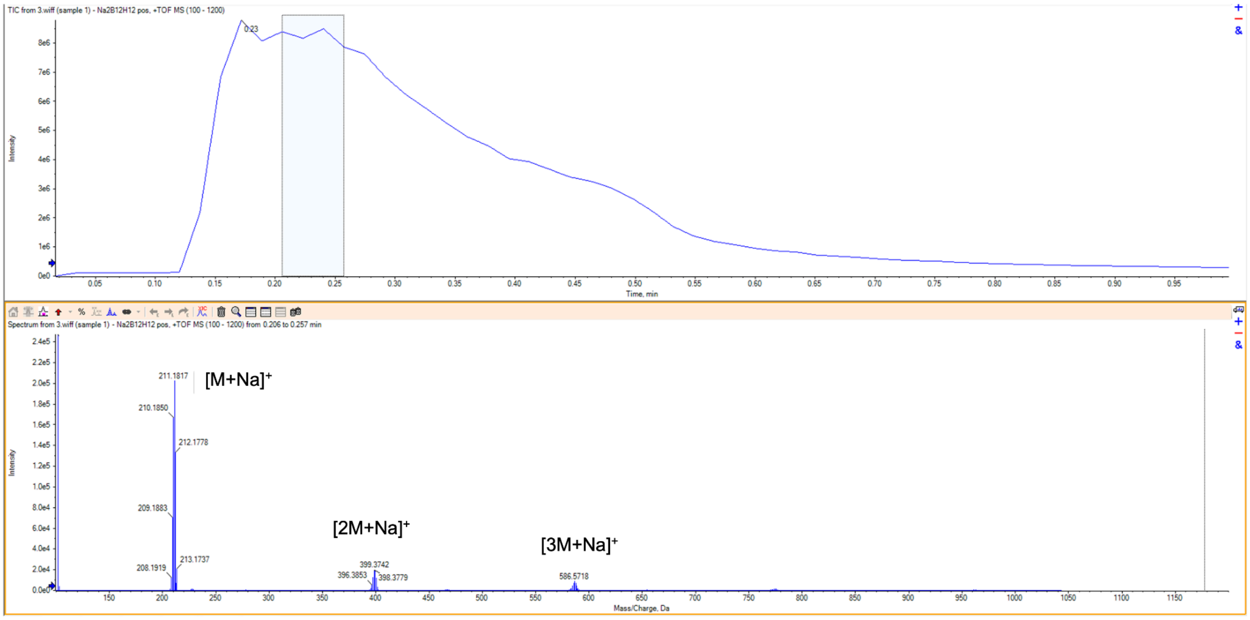


**Supp Fig 32. ESI-MS results of $Na_2B_{12}H_{12}$.** The total ion chromatogram and mass spectrum of $Na_2B_{12}H_{12}$ were shown.

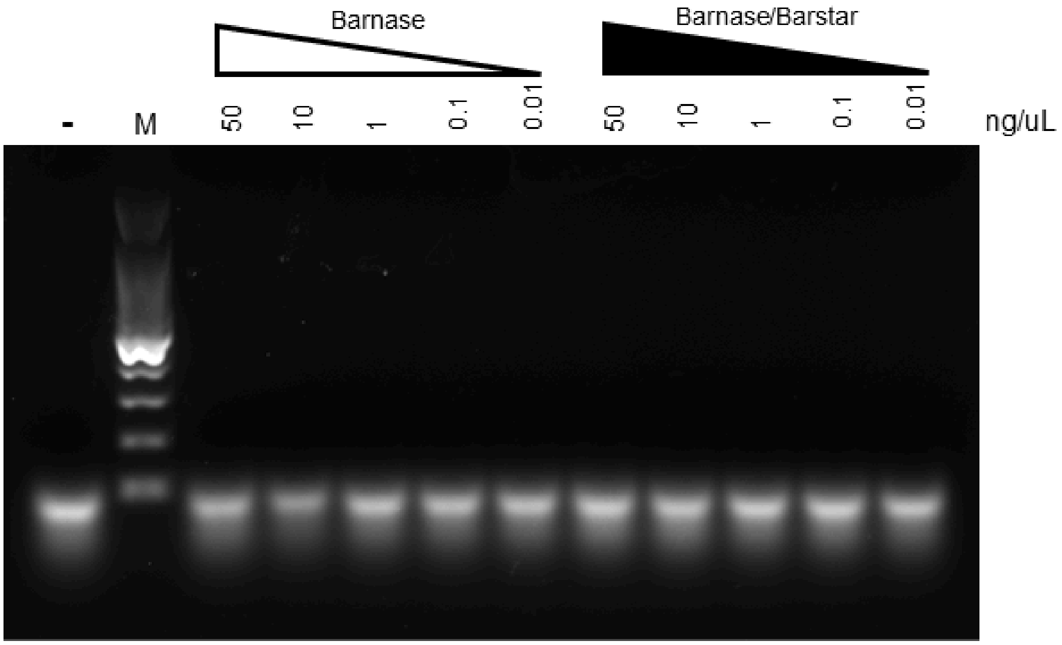


**Supp Fig 33. Attempted barnase activity assay** was performed in a serial dilution of free Barnase or Barnase/Barstar complex (1:1 in molar ratio) with ssRNA as the substrate. The dilution starting and ending concentrations of free Barnase or Barnase/Barstar complex were as labelled in ng/µL. - indicates negative control (nuclease-free), and M indicates DNA ladder marker.

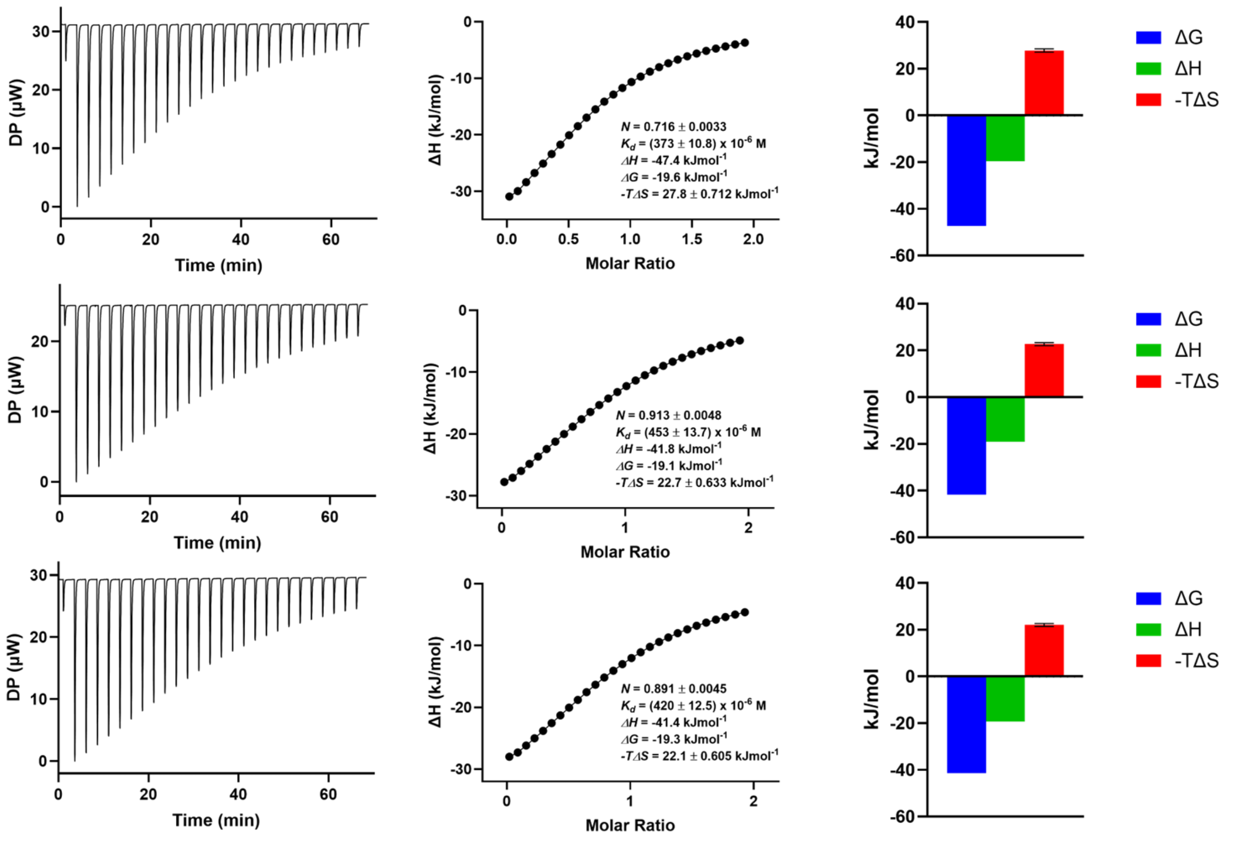


**Supp Fig 34. ITC results of $Na_2B_{12}H_{12}$ and 2HPγCD complexation in nuclease-free water.** Raw tracings, fitted heat plots and thermodynamic profilings of the titration experiments were shown. *N* = 3 replicates were performed.

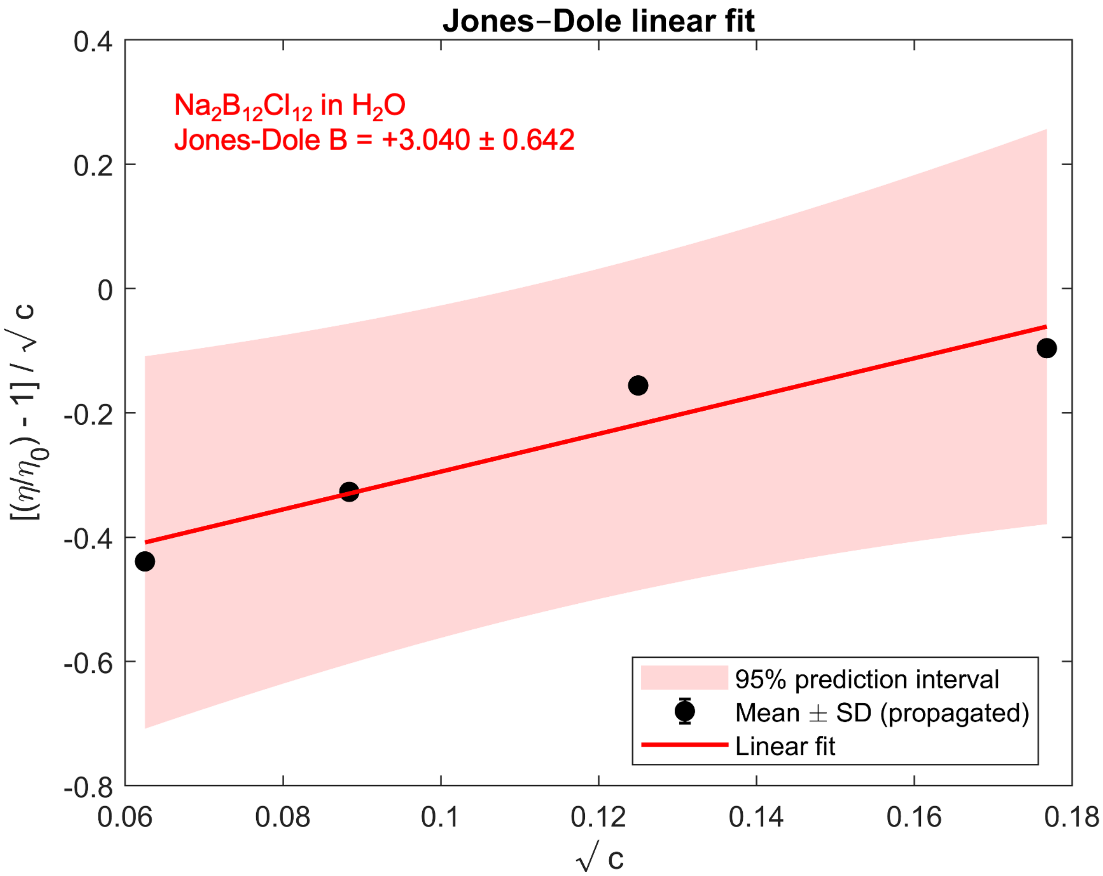


**Supp Fig 35. Solution dynamic viscosity changes with varying concentration of sodium dodecachloro-closo-dodecaborate, $Na_2[B_{12}Cl_{12}]$.** Related to **Fig. 6g**. The obtained Jones-Dole B value is +3.040 ± 0.642 (standard error of linear fit).

**Supp Table 1. ICP-OES results.**

| Element | Wavelength (nm) | Relative abundance (%) |
|---|---|---|
| B | 249.773 | 69.9 |
| Ca | 183.801 | 0.092 |
| K | 766.491 | 0.008 |
| Na | 589.592 | 24.1 |
| S | 182.034 | 0.065 |
| Si | 251.612 | 0.004 |
| Sn | 189.991 | 0.002 |
| Te | 238.578 | 0.013 |
| Tl | 190.864 | 0.007 |
| U | 409.014 | 0.009 |

**Supp Table 2.** ccfRNA sequencing reads and reference genome and gene mapping statistics.

| Sample | Total Clean Bases (Gb) | Clean Reads Q30 (%) | Clean Reads Ratio (%) | Total Reference Genome Mapping (%) | Total Gene Mapping Rate (%) |
|---|---|---|---|---|---|
| Subject 1, day 0, ctrl | 8.76 | 93.63 | 79.31 | 85.45 | 18.46 |
| Subject 1, day 0, $[B_{12}H_{12}]^{2-}$ | 8.83 | 93.15 | 75.62 | 84.62 | 16.03 |
| Subject 2, day 0, ctrl | 8.82 | 93.67 | 77.99 | 85.52 | 20.52 |
| Subject 2, day 0, $[B_{12}H_{12}]^{2-}$ | 8.83 | 93.59 | 76.09 | 76.88 | 17.39 |
| Subject 1, day 7, ctrl | 8.85 | 94.45 | 78.03 | 82.40 | 15.02 |
| Subject 1, day 7, $[B_{12}H_{12}]^{2-}$ | 8.84 | 94.03 | 69.58 | 81.23 | 10.45 |
| Subject 2, day 7, ctrl | 8.83 | 93.86 | 74.49 | 79.85 | 14.35 |
| Subject 2, day 7, $[B_{12}H_{12}]^{2-}$ | 8.83 | 94.22 | 66.72 | 80.29 | 24.68 |

**Supp Table 3**. Primers used in this study.

| # | Purpose | Strand | Sequence (5' → 3') | Amplicon size (bp) | Accession no. |
|---|---|---|---|---|---|
| 1 | Detects lambda DNA in human plasma | F | GCAAGTATCGTTTCCACCGT | 100 | LC730321.1 |
| 2 | Detects lambda DNA in human plasma | R | TTATAAGTCTAATGAAGACAAATCCC | | |
| 3 | Detects bacteriophage MS2 in human plasma | F | CGGCTGCTCGCGGATA | 65 | LC710217.1 |
| 4 | Detects bacteriophage MS2 in human plasma | R | AACTTGCGTTCTCGAGCGAT | | |
